# HAP40 functions as a proteostasis regulator by controlling huntingtin interactions and its release into the extracellular space

**DOI:** 10.64898/2026.03.31.715351

**Authors:** Eduardo Silva Ramos, Annett Boeddrich, Christian Haenig, Philipp Trepte, Orchid Ammar, Christopher Secker, Oliver Popp, Andranik Ivanov, Brandon Keith, Rachel J. Harding, Franziska Schindler, Tomas Koudelka, Leonard Roth, Nadine Scharek, Martina Zenkner, Nancy Neuendorf, Sabrina Golusik, Stephanie Beetz, Philipp Mertins, Ilaria Piazza, Sigrid Schnoegl, Erich E. Wanker

## Abstract

Huntingtin-associated protein 40 (HAP40) forms a stable protein complex with huntingtin (HTT). Its cellular function and how HAP40 loss influences mutant HTT (mHTT) abundance and pathobiology are currently unclear. Here, using diverse cellular models and OMICs methods, we demonstrate that HAP40 is an obligate interaction partner of full-length HTT and through its binding controls the abundance of HTT-associated proteins, indicating that it functions as HTT interaction regulatory unit. Also, loss of HAP40 in mHTT-expressing striatal cells impairs autophagosome-lysosome flux, triggers massive transcriptional dysregulation, including the activation of the CLEAR network, demonstrating that it functions as proteostasis regulator that acts on quality control pathways. Finally, mHTT-expressing cells lacking HAP40 showed increased secretion of mHTT through the ER-to-Golgi route, indicating that striatal cells reduce intracellular mHTT-induced proteotoxicity through activation of secretory pathways. Together, these results establish HAP40 as a critical proteostasis regulator that through controlling HTT interactions maintains cellular homeostasis.

## Introduction

Huntington’s disease (HD) is a monogenic, neurodegenerative disorder characterized by a triad of motor, cognitive and psychiatric symptoms(1, 2). It is caused by a CAG repeat expansion located in exon 1 of the *huntingtin* (*HTT*) gene, leading to the production of HTT protein with an abnormally elongated polyglutamine (polyQ) tract of largely unknown molecular function(1, 3). PolyQ-expanded HTT is believed to cause neurotoxicity through a dominant gain-of-function mechanism, potentially disrupting a wide range of cellular processes, including autophagy(4), transcription(5), splicing(6) and synaptic transmission(7). The pathogenic polyQ tract may also alter HTT’s normal physiological function in medium spiny neurons of the striatum, suggesting that both loss- and gain-of-function mechanisms may drive the pathology of HD. How the pathogenic polyQ tract in HTT causes dysfunction and selective neurodegeneration in HD brains is currently not well understood(8).

HTT is a large ∼350 kDa protein that is widely expressed in different cell-types and tissues(3). In mice, HTT was shown to be essential for early embryogenesis(9), while its ablation in neurons of adult animals was non-deleterious(10). Full-length HTT is largely composed of α-helical “HEAT” repeats that form folded super-helical structures(11) and are thought to function as protein-protein interaction (PPI) modules(12). In line with this, thousands of HTT-associated proteins have been identified in previous interaction screens with full-length HTT or its fragments(13–15), suggesting that HTT functions as an interaction hub in cells and directly binds diverse partner proteins involved in different subcellular processes(3). However, despite our knowledge about HTT PPIs, it remains largely unclear through which domains HTT interacts with partner proteins and how full-length HTT potentially controls the function of associated cellular proteins. Also, it is unknown how interaction partners specifically influence HTT function and control its abundance in mammalian cells. Knowledge about the latter is crucial for the development of improved HTT lowering strategies, because proteins that regulate mHTT abundance may be interesting targets for therapeutic development.

The ∼40 kDa HTT-associated protein (HAP40; F8A1) stands out among the ∼3,000 previously reported human HTT interaction partners(14, 16). It binds to full-length HTT in a cleft and stabilizes its secondary structure(16). Through the association of HAP40, a compact heterooligomeric HTT-HAP40 complex is formed that likely interacts with other proteins and thereby may regulate their specific activities. Recent structural and functional investigations with recombinant proteins, cell models and *Drosophila melanogaster* support the hypothesis that HTT-HAP40 heterooligomers are functionally relevant(17, 18). This is also supported by overexpression and gene knock-down experiments, indicating that HTT overproduction can increase HAP40 protein levels, while HTT depletion dramatically decreases HAP40 abundance in cells(19, 20).

Currently, the cellular function of HAP40 is unknown. Also, it is unclear how HAP40 influences mHTT abundance and pathobiology in mammalian cells. Experimental evidence has demonstrated that full-length HTT interacts with the cargo receptor p62 and functions as a scaffold for selective autophagy(21, 22), suggesting that binding of HAP40 to HTT might influence this protein degradation process in cells. Furthermore, it was shown that HTT directly associates with the Golgi apparatus(23, 24) and facilitates vesicle trafficking to the plasma membrane(25), suggesting that this subcellular process may also be influenced by HAP40. An association of HAP40 with the GTPase Rab5, which regulates vesicle fusion and cargo delivery to endosomes and lysosomes(26), has also been described(27). Whether inactivation of HAP40 influences specific protein degradation pathways or vesicle transport processes, however, remains unclear.

In this study, we comprehensively assessed the cellular function of HAP40, utilizing multiple HAP40 knock-out (KO) cell lines, OMICs methods as well as a set of biochemical and functional assays. Also, we addressed the question of how HAP40 controls mHTT abundance and pathobiology in mammalian cells. We found that HAP40 is an obligate HTT interaction partner and controls its association with other cellular proteins, indicating that it functions as a HTT interaction regulatory unit that indirectly controls the abundance and function of HTT-associated proteins. Also, we observed that depletion of HAP40 in mHTT-expressing mouse striatal ST*Hdh*^Q111^ cells causes massive transcriptional dysregulation, activation of the CLEAR (Coordinated Lysosomal

Expression and Regulation) pathway and impairment of autophagy, indicating that HAP40 functions as a critical proteostasis regulator, which controls the activity of protein degradation pathways. Finally, we found that in striatal ST*Hdh*^Q111^-*Hap40*KO cells the secretion of full-length mHTT into the extracellular space is significantly increased, indicating that cells can reduce mHTT abundance and proteotoxicity through activation of secretory pathways. The potential implications of our findings for the development of novel causal therapies for the treatment of HD are discussed.

## Results

### Loss of HAP40 decreases the abundance of full-length HTT in HEK293 and striatal cell lines

Previous investigations suggest that HAP40 and HTT protein levels under physiological conditions are interdependent(17). To further explore this relationship, we used CRISPR/Cas9 genome editing to generate a HEK293 HAP40 knockout (HEK293-HAP40KO) cell line (**Fig. S1A**). We found that HTT^Q23^ protein levels are ∼50% lower in HEK293-HAP40KO cells than in wild-type (WT) cells (**Fig. 1A and B**). In comparison, the depletion of HTT^Q23^ in HEK293-HTTKO cells caused an almost complete loss of HAP40 (**Fig. 1A and B**), confirming previous reports(17). The levels of both HAP40 and HTT^Q23^ were rescued at least in part, when the respective proteins were overproduced in HAP40KO or HTTKO cell lines (**Fig. S1B and C**).

**Fig. 1.**
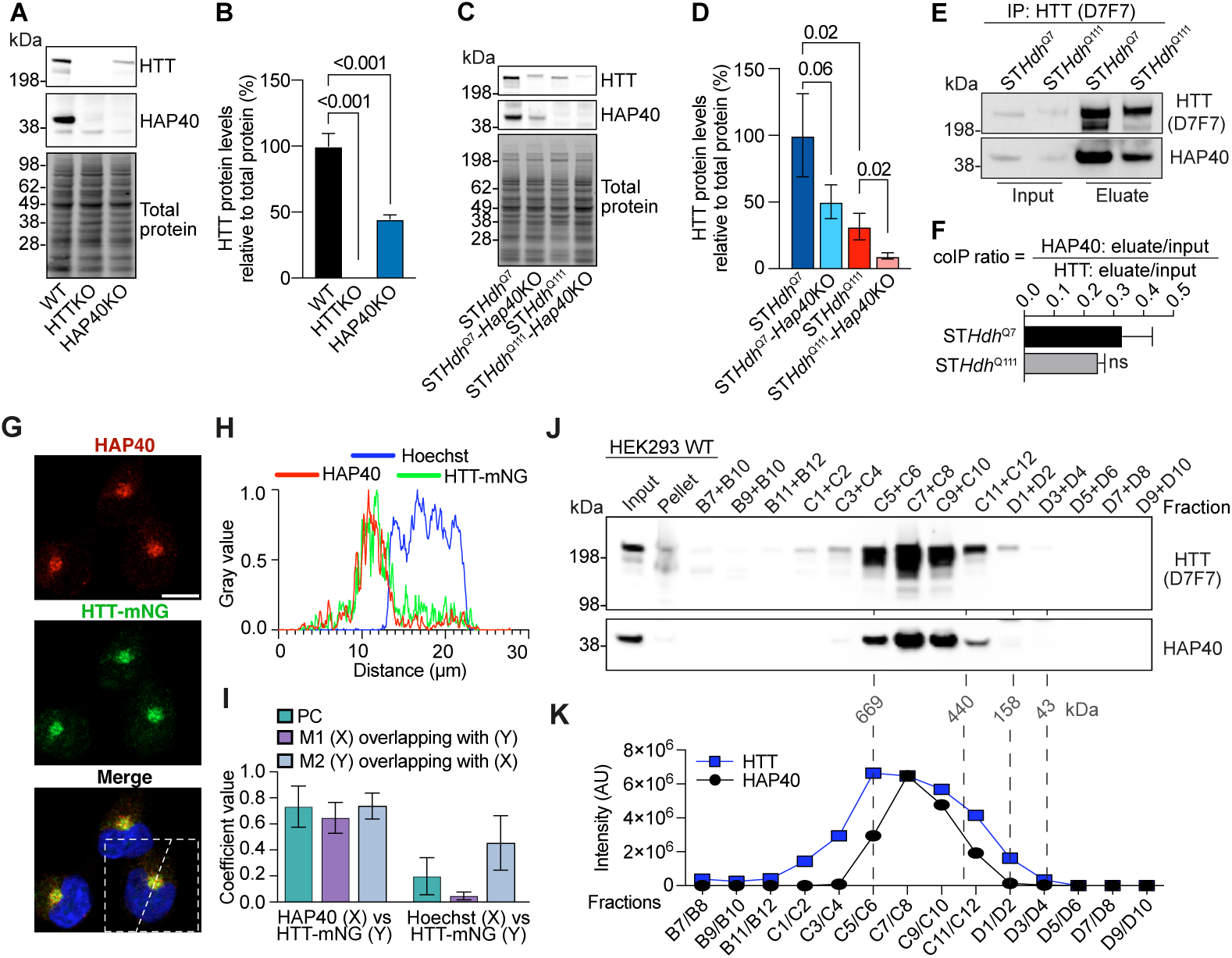
HTT and HAP40 form a stable complex. **A)** Representative immunoblot of HEK293 total cell lysates. Membranes were probed with antibodies against HTT (anti-HTT D7F7) and HAP40. Total protein was visualized using No-Stain protein labelling reagent and served as a loading control. **B)** Quantification of HTT protein levels in **A**, expressed relative to total protein and normalized to HEK WT. Data are presented as mean ± SD of four biological replicates. Statistical significance was determined via one-way ANOVA followed by Dunnett’s multiple comparisons test. **C)** Representative immunoblot of different mouse striatal total cell lysates. Membranes were probed with antibodies against HTT (anti-HTT D7F7) and HAP40. Total protein was visualized using No-Stain protein labelling reagent and served as a loading control. **D)** Quantification of HTT protein levels in **C**, expressed relative to total protein and normalized to ST*Hdh*^Q7^. Data are presented as mean ± SD of three biological replicates. Statistical significance was determined using a Student’s t-test. **E)** Representative immunoblots of immunoprecipitations of HTT with anti-HTT (D7F7) antibody using cultured mouse striatal cell lysates. IP fractions represent 5% input sample and 100% eluate. Samples were subjected to SDS-PAGE and immunoblotted for HTT using anti-HTT (D7F7) and anti-HAP40. **F)** Quantification of the co-IP enrichment ratio of HAP40 relative to HTT from mouse striatal HTT immunoprecipitations shown in **E**. Eluate band intensities for HAP40 and HTT were normalized to their respective input band intensities. The enrichment value of HAP40 was then normalized to that of HTT. Data are presented as mean ± SD of three biological replicates. No significant difference (ns) was observed (Student’s t-test). **G)** Representative confocal images of HEK293-HTT-mNG cells immunostained for HAP40 (red), mNeonGreen (green), with nuclei counterstained using Hoechst (blue). The merged image includes an inset showing the line used for line-scan analysis. Images were acquired at 63x magnification. Scale bar: 10 µm. **H)** Line-scan analysis of cell shown in **G**, displaying fluorescence intensity (gray values) from all three channels plotted against distance across the line. **I)** Bar graph showing colocalization analysis of HAP40 (X) and HTT-mNG (Y) signals, as well as Hoechst (X) and HTT-mNG (Y) as a negative control. The analysis includes the average Pearson’s correlation coefficient (PC) and Manders coefficients M1 (fraction of X overlapping with Y) and M2 (fraction of Y overlapping with X). Data represent mean ± SD of three biological replicates. **J)** Representative size-exclusion chromatography of HEK293 cell total lysates. Fractions were pooled by combining two wells from a 96-well plate, yielding 14 final fractions. Proteins were acetone-precipitated and sequentially subjected to SDS-PAGE followed by immunoblotting using anti-HTT (D7F7) and anti-HAP40 antibodies. From input and pellet samples 10 µg were loaded, while 10% of each fraction pool was loaded. Molecular weight markers of size-exclusion control proteins are displayed below the HAP40 immunoblot. Two independent experiments were performed. **K)** HTT and HAP40 immunoblot intensities in fractions B7 to D10 of SEC from **J** were quantified using iBright Analysis Software. On top of the diagram molecular weight standards are shown which were used for calibration of the SEC column. Blue line with squares reveals HTT intensities, black line with circles reveals HAP40 intensities. Band intensities are shown as an arbitrary value (AU) and plotted against the pooled fractions.

To assess whether these effects extend to mouse striatal cells and pathogenic HTT alleles, we used homozygous cell lines(28) that endogenously express either non-pathogenic (ST*Hdh*^Q7/Q7^; here on ST*Hdh*^Q7^) or pathogenic (ST*Hdh*^Q111/Q111^; here on ST*Hdh*^Q111^) full-length HTT. We generated *Hap40*KO derivatives of both lines using CRISPR/Cas9 (**Fig. S1D and E**). In the unmodified parental lines, we found that steady-state HTT^Q111^ protein levels are ∼50% lower than HTT^Q7^ levels (**Fig. 1C and D**), in line with previous observations(29, 30). Deletion of HAP40 further reduced HTT protein levels in both ST*Hdh*^Q111^-*Hap40*KO and ST*Hdh*^Q7^-*Hap40*KO cell lines (**Fig. 1C and D**), indicating that loss of HAP40 lowers both wild-type and mutant HTT protein abundances in striatal cells. To assess whether transcriptional dysregulation is responsible for the HTT protein abundance changes, we quantified HTT transcript levels utilizing an established q-PCR assay(31). This analysis revealed that both *Htt*^Q111^ and *Htt*^Q7^ transcript levels were mildly decreased in striatal *Hap40*KO cells compared to controls (**Fig. S1F**), suggesting that transcriptional changes do not account for the observed ∼50% reductions in protein levels.

### HAP40 is an obligate interaction partner of full-length HTT

Studies with recombinant human proteins have previously shown that HAP40 and full-length HTT form a stable 1:1 complex(16, 19). However, their interaction under physiological conditions remains less well characterized. To address this, we first performed co-immunoprecipitations (co-IPs) to enrich HTT-HAP40 complexes from crude protein extracts of striatal cells. The anti-HTT antibody D7F7 was utilized for co-IPs, because it recognizes an epitope of HTT not required for HAP40 binding, based on published HTT-HAP40 cryo-EM structural studies (**Fig. S1G**). We observed that similar amounts of HAP40 are enriched from striatal ST*Hdh*^Q7^ and ST*Hdh*^Q111^ cell extracts (**Fig. 1E and F**), indicating that both HTT^Q7^ and HTT^Q111^ bind similar amounts of HAP40 under physiological conditions. A similar result was also obtained when HTT-HAP40 complexes were enriched from crude brain extracts of zQ175 HD knock-in(32) and control mice using the D7F7 antibody (**Fig. S1H**).

Next, to investigate the co-localization of HTT and HAP40, we generated a HEK293 knock-in cell line expressing HTT^Q23^ fused at its C-terminus with mNeonGreen (HTT^Q23^-mNG) (**Fig. S1I**). Immunolabeling of HTT^Q23^-mNG producing cells with an anti-HAP40 antibody revealed a high degree of HTT^Q23^-mNG and HAP40 co-localization in distinct foci in the perinuclear region of HEK293 cells (**Fig. 1G-I**). This supports the hypothesis that both proteins form a complex under physiological conditions. Very similar results were obtained when wild-type and HTT^Q23^-mNG-producing HEK293 cells were immunolabeled with anti-HTT and anti-HAP40 antibodies (**Fig. S2A-H**). The specificities of the applied antibodies for their respective targets were confirmed with HEK293-HTTKO and HEK293-HAP40KO cell lines (**Fig. S2I and J**). Next, the localization of apo-HTT was examined in HEK293-HAP40KO cells using the validated anti-HTT antibody. In stark contrast to the perinuclear foci of HTT in WT cells, HEK293-HAP40KO cells displayed a weaker perinuclear localization and a more dispersed cellular distribution of apo-HTT (**Fig. S2K**).

Finally, we performed size-exclusion chromatography (SEC) experiments using HEK293 cells to determine the size distribution of endogenously formed HTT-HAP40 heterooligomers. Analysis of protein samples by SDS-PAGE and immunoblotting revealed molecular weights between ∼440 and ∼670 kDa for the size-fractionated HTT^Q23^ and HAP40 proteins (**Fig. 1J, K and Fig. S3A, B**), indicating that complexes formed under physiological conditions may be larger than the reported 1:1 ∼390 kDa HTT-HAP40 heterooligomers(16). This suggests that additional proteins may be bound to the HTT-HAP40 complex in HEK293 cells. Interestingly, perfectly overlapping elution profiles of HAP40 and HTT^Q23^ were obtained (**Fig. 1K**), indicating that most if not all HAP40 protein is stably bound to full-length HTT^Q23^ in HEK293 cells. In comparison, HTT^Q23^, but not HAP40, was detected in high molecular weight protein fractions (>669 kDa), suggesting that at least a small amount of endogenously produced HTT^Q23^ is not bound by HAP40 in cells. Moreover, an analysis of mouse brain extracts with SEC yielded similar co-fraction profiles (**Fig. S3C**), confirming that HAP40 is an obligate interaction partner of full-length HTT under native conditions.

### HAP40 binding alters the conformation of full-length HTT^Q23^ in mammalian cells

We hypothesized that loss of HAP40 may alter the structure of full-length HTT and perturb its protein-binding properties. To address this, we first investigated the interaction between HTT^Q23^ and HAP40 utilizing a well-established quantitative BRET assay(33). With this method the interactions between nanoluciferase (NL, donor) and ProteinA-mCitrine (PA-mCit, acceptor)-tagged fusion proteins (e.g., NL-protein#1 and PA-mCit-protein#2) can be detected with high specificity and sensitivity in cells, when the sensor proteins NL and PA-mCit are in close proximity (<10 nm). The available cryo-EM structures of HTT-HAP40 heterooligomers(16, 19) suggest that both the N- and C-termini of HTT and HAP40 are in close proximity (**Fig. S4A**), suggesting that N-N and C-C tagging configurations with sensor proteins might reveal BRET, while N-C and C-N tagging might not, because the sensor proteins are far apart. Strikingly, our measurements revealed strong BRET signals, when N-N (NL-HTT^Q23^ vs PA-mCit-HAP40) or C-C (HTT^Q23^-NL vs HAP40-mCit-PA) tagged reporter protein fusions were co-produced in HEK293 wild-type cells (**Fig. 2A and B**). In contrast, BRET signals were undetectable with the N-C (NL-HTT^Q23^ vs HAP40-mCit-PA) or C-N (HTT^Q23^-NL vs PA-mCit-HAP40) tagging configurations, confirming our structure-based predictions.

**Fig. 2.**
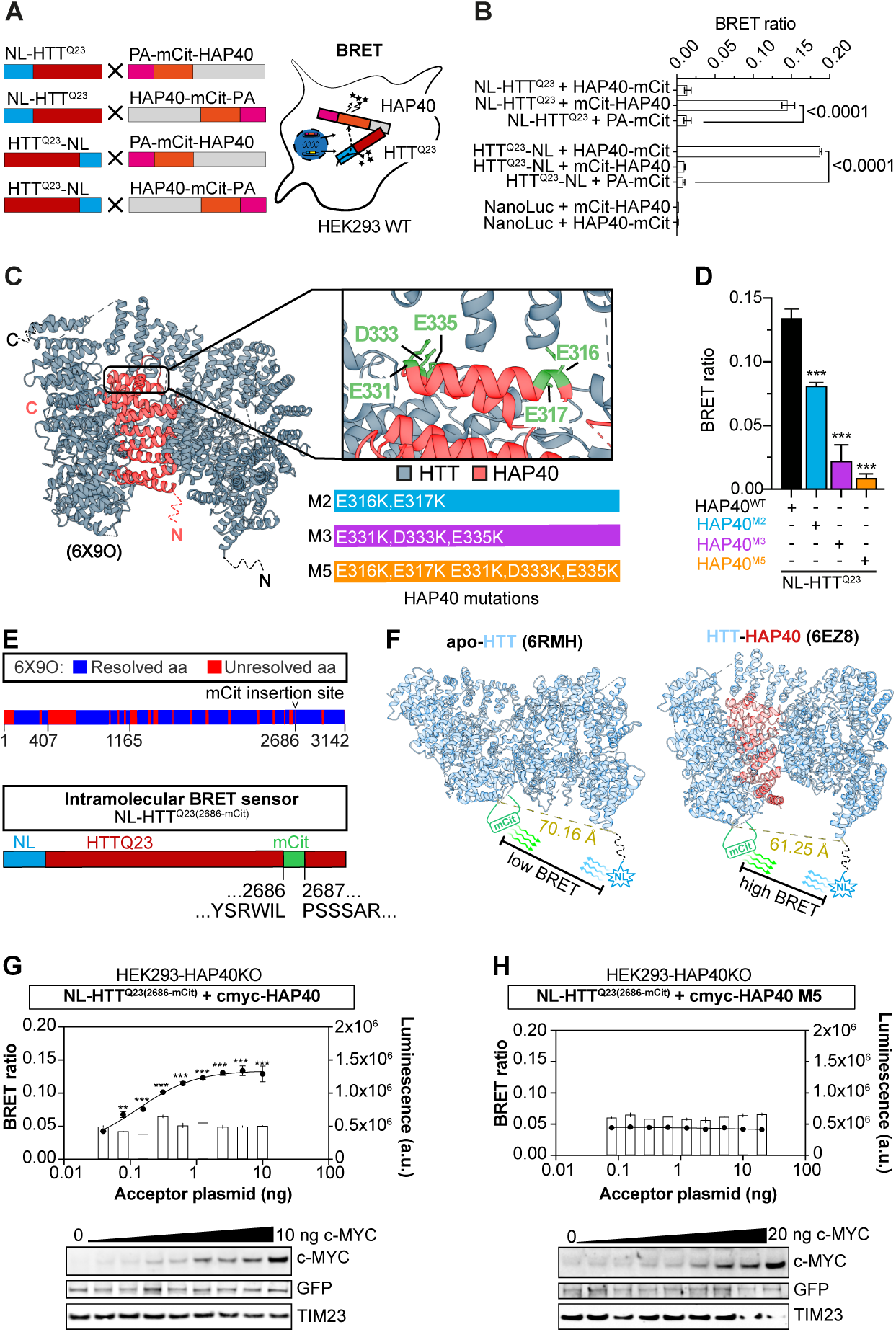
HAP40 residues E316, E317, E331, D333 and E335 are required for the formation of a stable HTT-HAP40 complex. **A)** Schematic representation of the BRET assay workflow. Expression vectors encoding c-myc-NL and PA-mCit tagged proteins were co-transfected into HEK293 cells. Binary interactions were detected in vivo using in-cell bioluminescence resonance energy transfer (BRET). **B)** BRET ratios of binary interactions between NanoLuc- and mCitrine-tagged full-length HTT^Q23^, HAP40, and respective controls. Data are presented as mean ± SD (n=2), each with three technical replicates. Statistical significance was determined via two-way ANOVA followed by Bonferroni’s multiple-comparisons test. **C)** Left: Cryo-EM structure of HTT-HAP40 (6X9O). HAP40 is shown in red and HTT in blue. Right: Zoomed-in view displaying the five conserved HAP40 amino acid residues (gold: E316, E317, E331, D333, E335) and the three HAP40 mutant variants generated based on the number of amino acids converted into lysine. **D)** BRET ratios of binary interactions between c-myc-NL-HTT^Q23^ and PA-mCit-HAP40 (wild-type and mutants). Data are presented as mean ± SD, n=2, each with three technical replicates. Statistical significance was determined via two-way ANOVA followed by Dunnett’s multiple-comparisons test. **E)** Top: Linear schematic representation of resolved (blue) and unresolved (red) segments of the Cryo-EM structure of HTT-HAP40 (6X9O), indicating the mCitrine (mCit) insertion site. Below: schematic representation of the HTT intramolecular BRET sensor (NL-HTT^Q23(2686-mCit)^), displaying the tagging positions of NanoLuc at the N-terminus and mCitrine between amino acids 2686 and 2687 of human HTT. **F)** Cryo-EM structures of apo-HTT (6RMH) and HTT-HAP40 (6EZ8), illustrating the distances between resolved residues near the mCitrine insertion site and the N-terminus of HTT. Distances are shown in angstroms (Å). The relative positions of NanoLuc and mCitrine are indicated, along with predicted differences in BRET signal based on donor-acceptor distance. **G)** BRET saturation assay with co-transfection of the HTT intramolecular BRET sensor (150 ng, pcDNA-NL-HTT^Q23(2686-mCit)^) and increasing amounts of pcDNA-c-myc-HAP40 (wild-type; 0-10 ng) in HEK293-HAP40KO cells. The plot displays BRET ratio values (dashed line) and luminescence values (bar graph). Data are presented as mean ± SD (n=2). Statistical significance was determined via one-way ANOVA followed by Dunnett’s multiple-comparisons test. Corresponding immunoblots related to BRET assay using anti-c-myc, anti-GFP, and anti-TIM23 (loading control) antibodies are shown below. **H)** BRET saturation assay using the HTT intramolecular BRET sensor with increasing amounts of c-myc-HAP40 (5M mutant). Experimental conditions and statistical analysis were performed as described in **G**.

Next, we assessed whether BRET measurements are sensitive enough to detect the impact of amino acid changes on HTT-HAP40 interactions. Informed by the structure of the HTT-HAP40 complex, we focused on five negatively charged residues in HAP40 (E316, E317, E331, D333 and E335), which are predicted to be critical for complex stability(16). We generated three PA-mCit-tagged HAP40 protein variants in which two (E316K, E317K; M2), three (E331K, D333K and E335K; M3) or five (E316K, E317K, E331K, D333K and E335K; M5) negatively charged residues were exchanged for positive lysine residues (**Fig. 2C**). Then, each of these proteins was co-produced together with NL-HTT^Q23^ in HEK293 cells and BRET ratios were quantified. In comparison to the wild-type protein (PA-mCit-HAP40^WT^) the protein variants PA-mCit-HAP40^M2, M3^ ^and^ ^M5^ showed significantly reduced BRET values (**Fig. 2D**). As expected, the BRET signal was lowest with the PA-mCit-HAP40^M5^ protein, which bears the largest surface charge change.

Finally, to investigate the impact of amino acid exchanges on the stability of HTT-HAP40 heterooligomers, we co-produced each protein variant together with HTT^Q23^-FLAG in insect cells and co-purified protein complexes with FLAG-affinity and size-exclusion chromatography(19). We observed protein complexes with HAP40^WT^ and all investigated protein variants (HAP40^M2, M3^ ^and^ ^M5^). Compared to HAP40^WT, M2^ ^and^ ^M3^, however, co-enrichment of HAP40^M5^ was low (**Fig. S4B**), indicating that this variant of HAP40 binds weakly to HTT^Q23^. This result is also supported by differential scanning fluorimetry experiments (**Fig. S4C**), where a gradual decrease in stability is observed with increasing mutations, further indicating that in comparison to purified HTT^Q23^-HAP40^WT^ heterooligomers, HTT^Q23^-HAP40^M5^ complexes have a much lower thermal stability.

Based on the reported cryo-EM structures(16, 19, 34), we hypothesized that full-length HTT^Q23^ in the presence and absence of HAP40 in cells might be structurally distinct. To test this in live cells, we engineered the intramolecular BRET sensor NL-HTT^Q23(2686-mCit)^ (**Fig. 2E**). In this protein NL is fused to the N-terminus of HTT and mCitrine is incorporated into an unstructured loop at leucine 2686. We reasoned that HAP40 binding might induce conformational compaction of HTT^Q23^, bringing the N- and C-terminal regions into closer proximity and thereby increasing the BRET signal in cells (**Fig. 2F**). Consistent with this, co-production of NL-HTT^Q23(2686-mCit)^ together with HAP40^WT^ in HEK293-HAP40KO cells revealed a significant BRET increase (**Fig. 2G**), while such an effect was not observed with the HAP40^M5^ protein (**Fig. 2H**), which binds very weakly to HTT^Q23^ in cells (**Fig. 2D**). This indicates that the molecular distance between the N-terminus and the unstructured C-terminal loop region (L2686) is significantly larger in HTT^Q23^ in the absence than in the presence of HAP40, confirming previously reported structural models(19). Importantly, SDS-PAGE and immunoblot analyses confirmed that both HAP40^WT^ and HAP40^M5^ were expressed at comparable levels when co-produced with the BRET sensor in HEK293-HAP40KO cells (**Fig. 2G and 2H**), ruling out expression differences as a confounding factor.

## Loss of HAP40 alters HTT’s interaction profile

We hypothesized that binding of HAP40 to HTT^Q23^ might influence its association with other proteins in cells. To uncover HTT^Q23^ interaction partners, we employed an immunoprecipitation-mass spectrometry (IP-MS) approach, which enables the detection of PPIs with high specificity and sensitivity using label-free quantification(35). A pilot experiment was performed with the anti-HTT antibody D7F7, demonstrating efficient enrichment of the HTT-HAP40 complex from crude WT protein extracts (**Fig. 3A**). Building on this, we performed more comprehensive IP-MS experiments with extracts from HEK293 WT, HEK293-HTTKO and HEK293-HAP40KO cells (**Fig. 3B)**. Given our earlier findings that indicate HAP40 is an obligate HTT interactor (**Fig. 1J**), we anticipated that D7F7 would predominantly co-enrich HTT-HAP40-associated proteins from WT extracts, while apo-HTT^Q23^ associated proteins are co-enriched from HAP40KO protein extracts. We used HEK293-HTTKO extracts to evaluate nonspecific antibody binding (**Fig. 3B**) and quantified relative protein abundances in the antibody-enriched samples using intensity-based absolute quantification (iBAQ) (**Table S1**).

**Fig. 3.**
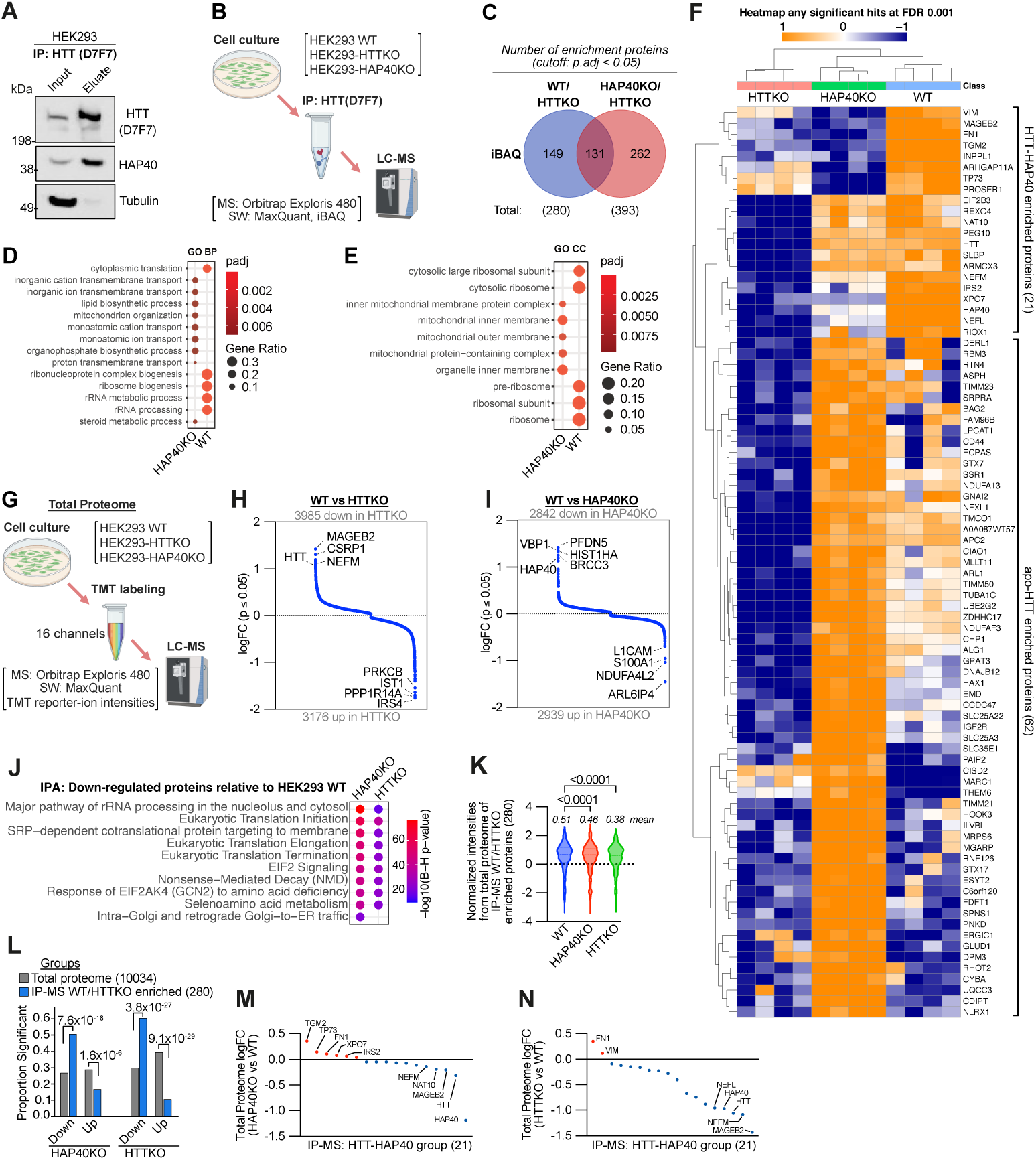
HTT interactome changes in the absence of HAP40. **A)** Immunoblots of HTT immunoprecipitations from wild-type HEK293 lysates using anti-HTT D7F7. Lanes display 5% input and 80% eluate relative to starting material. Samples were analyzed by SDS-PAGE and probed for HTT and HAP40. Data are representative of four independent experiments. **B)** Schematic of the HTT IP-MS workflow. Three HEK293 cell lines (wild-type, HTTKO, and HAP40KO) were subjected to immunoprecipitation with anti-HTT (D7F7) followed by LC-MS/MS analysis. Four biological replicates of each cell line were used. **C)** Quantification of proteins significantly enriched (adjusted p < 0.05) by co-IP in the two experimental contexts: (i) HTT-HAP40 complexes isolated from wild-type cells and (ii) apo-HTT complexes isolated from HAP40KO cells. Proteins enriched in HTTKO extracts were excluded. The Venn diagram illustrates the overlap between the two groups, with the intersecting region shown at the center. **D)** Gene Ontology (GO) enrichment analysis of biological process (BP) terms and **E)** cellular component (CC) terms for proteins significantly enriched from HTT-IPs in wild-type HEK293 cells or in HAP40KO cells (apo-HTT^Q23^), excluding proteins also enriched in HTTKO extracts. Terms were identified using the enrichGO package in R and ranked by Benjamini–Hochberg FDR-adjusted p-values. **F)** Heatmap of iBAQ intensities (row-wise Z-score normalized) for proteins that passed a stringent enrichment threshold (FDR ≤ 0.001) across the three HEK293 genotypes. Hierarchical clustering was performed using Euclidean distance, revealing distinct enrichment patterns among the cell lines. **G)** Overview of the HEK293 total proteome workflow. Three HEK293 cell lines (wild-type, HTTKO, and HAP40KO) were processed for Tandem Mass Tag (TMT-16-plex) labeling and subsequent LC-MS/MS analysis. Four independent biological replicates were prepared for each genotype. **H)** Scatter plot of protein-level changes between HEK293 wild-type and HTTKO cells (total proteome TMT-MS). The vertical axis shows the log fold change (logFC) of each protein, a significance cutoff of adjusted p < 0.05 (Benjamini–Hochberg FDR correction) is indicated. Positive values denote higher abundance in wild-type, negative values denote higher abundance in the HTTKO. **I)** Same representation as in **H** but contrasting wild-type with HAP40KO cells. Axes and statistical thresholds are identical. **J)** Ingenuity Pathway Analysis (IPA) of proteins significantly downregulated (adjusted p ≤ 0.05) in both HAP40KO and HTTKO versus wild-type HEK293 cells. Canonical pathways are ranked by Benjamini–Hochberg FDR-adjusted p-values. **K)** Cross-reference of IP-MS hits (280 WT/HTTKO enriched proteins) against the total proteome TMT-MS dataset. Normalized intensities from the three HEK293 cell lines were used for comparison. **L)** Enrichment analysis of differentially regulated proteins. Two groups were examined: (i) the 280 IP-MS WT/HTTKO enriched proteins and (ii) the global HEK293 TMT-MS proteome in either HAP40KO or HTTKO cells. For each group, proteins were classified as significant or non-significant (adjusted p ≤ 0.05) based on the TMT-MS data. Fisher’s exact test was applied to assess enrichment; p-values are displayed above the bars. **M)** LogFC values for the 21 proteins that clustered together in the hierarchical analysis shown in **F** (HTT-HAP40 enriched protein set) plotted against the total proteome TMT-MS data for HAP40KO versus wild-type. **N)** Same 21-protein set as in **M**, now plotted for the HTTKO versus wild-type comparison. All statistical analyses employed Benjamini–Hochberg correction for multiple testing unless otherwise noted. **B)** and **G)** were created in BioRender. Wanker, E. (2026) https://BioRender.com/ci9qwne.

Principal component analysis (PCA) revealed that protein abundance profiles from D7F7-enriched samples clustered well within biological replicates (**Fig. S5A**). Furthermore, the PCA showed that sample groups display HAP40 dependency, indicating that HTT^Q23^ in the presence and absence of HAP40 associates with different sets of cellular proteins. To identify specific apo-HTT^Q23^ interactors, we evaluated protein enrichment by statistical significance (**Fig. 3C and Fig. S5B and C**). Only proteins that were reproducibly enriched (adj. p-value <0.05) with D7F7 from WT or HEK293-HAP40KO protein extracts but not from HEK293-HTTKO extracts were considered as HTT interaction partners (**Fig. S5B and C**). With these criteria, we detected a total 542 unique HTT^Q23^-associated proteins, of which 149 were exclusively co-enriched from WT and 262 from HEK293-HAP40KO protein extracts (**Fig. 3C**). Notably, 131 proteins (∼20%) were shared between both groups, indicating that the absence of HAP40 significantly alters HTT’s interaction landscape. Compared with the HTT^Q23^-HAP40 complex, apo-HTT^Q23^ in HEK293-HAP40KO cells bound a greater number of proteins, although it is significantly less abundant (**Fig. 1A and B**), suggesting that it is more interaction-prone and aberrantly associates with various cellular proteins (**Fig. 3C)**. Gene Ontology (GO) term enrichment analysis supported this conclusion. Apo-HTT^Q23^ associated with a diverse array of functionally distinct proteins, including those involved in mitochondrial processes, intracellular transport, and lipid biosynthesis (**Fig. 3D and E**). In strong contrast, proteins involved in ribosome biogenesis and rRNA metabolism were predominately co-enriched with the HTT^Q23^-HAP40 complex **(Fig. 3D and E)**. KEGG pathway enrichment analysis yielded similar results (**Fig. S5D and E**). Cluster analysis of high-confidence interactors (adj. p-value <0.001) confirmed two distinct groups: one preferentially associated with HTT^Q23^-HAP40 and the other with apo-HTT^Q23^ **(Fig. 3F and Fig. S5F),** whereby the apo-HTT^Q23^ group was larger. GO term enrichment analysis revealed that apo-HTT^Q23^-bound proteins were involved in mitochondrial and transport functions, whereas the HTT^Q23^-HAP40-bound proteins were involved in filament assembly and trafficking pathways (**Fig. S5G**). Collectively, these data suggest that under physiological conditions, HTT^Q23^, in complex with HAP40, preferentially associates with proteins involved in cytoplasmic translation, ribosome biogenesis and vesicle-cytoskeletal trafficking, while in the absence of HAP40 it binds to proteins involved in dispersed subcellular processes. To validate these findings, we used a quantitative cell-based binding assay to assess the impact of HAP40 loss on HTT interactions. We focused on the E3 ubiquitin-protein ligase RNF126(36), which was enriched with apo-HTT^Q23^ but absent from HTT^Q23^-HAP40 fractions **(Fig. 3F)**. Using HEK293 WT and HEK293-HAP40KO cells, we co-produced full-length NL-HTT^Q23^ and PA-mCit-RNF126 and quantified *in-cell* BRET. The BRET signal was significantly higher in HEK293-HAP40 KO cells compared to WT cells, confirming that RNF126 preferentially interacts with HTT^Q23^ in the absence of HAP40 (**Fig. S5H**).

Finally, immunoprecipitations of HTT using striatal cells, demonstrated a similar shift in GO enrichment terms upon the loss of HAP40, where HTT-HAP40 enriched from ST*Hdh*^Q7^ or ST*Hdh*^Q111^ cells predominantly associated with proteins involved in ribosome biogenesis and translation, while apo-HTT enriched from ST*Hdh*^Q7^-*Hap40*KO or ST*Hdh*^Q111^-*Hap40*KO cells associated with other proteins (**Fig. S6A-C)**.

### Loss of HAP40 decreases the abundance of HTT-associated proteins in HEK293 cells

To assess global proteomic changes resulting from HAP40 or HTT knockout in HEK293 cells, we performed Tandem Mass Tag (TMT)-based mass spectrometry for total proteome analysis across HEK293 WT, HAP40KO, and HTTKO cell lines (**Fig. 3G, Table S2**). PCA of the TMT data revealed distinct clustering by genotype. HEK293-HAP40KO samples exhibited a moderate shift along both PCs, while HEK293-HTTKO samples displayed a substantial divergence (**Fig. S5I**). Comparative analysis revealed substantial alterations in the proteome of both KO lines relative to WT (**Fig. 3H and I)**. For example, HEK-HAP40KO cells showed ∼3,000 significantly downregulated proteins, while HEK-HTTKO cells exhibited a markedly larger effect, with ∼4,000 proteins downregulated (adjusted p-value ≤ 0.05; **Fig. 3H and I**). These data underscore the profound proteome-wide consequences of losing either HAP40 or HTT. To explore the functional relevance of these changes, we performed Ingenuity Pathway Analysis (IPA) on the significantly downregulated proteins in each KO line. Interestingly, in both HEK293-HTTKO and HEK293-HAP40KO cells proteins involved in rRNA processing and eukaryotic translation were significantly reduced in abundance (**Fig. 3J**), supporting the hypothesis from interaction studies that HTT and HAP40 may play a functional role in these core cellular processes (**Fig. 3D and E**).

We next examined the abundance of proteins identified as HTT^Q23^-HAP40 interactors in our IP-MS experiments. TMT intensity data revealed a significant reduction in their abundance in both KO lines (**Fig. 3K**). To determine whether this subset of proteins was disproportionately decreased in abundance, we performed Fisher’s exact tests comparing the proportion of significantly downregulated proteins in the interactor group versus the background proteome. In HEK293-HAP40KO cells, ∼50% of the HTT^Q23^-HAP40-associated proteins were significantly decreased in abundance, compared to 27% in the background proteome (Fisher’s exact test, p < 0.05). Similarly, in HEK293-HTTKO cells, ∼60% of the subgroup was significantly reduced in abundance, versus ∼30% of the background, indicating a significant proportion of proteins with reduced abundance among the HTT-HAP40-associated protein group (**Fig. 3L**). We observed similar trends among apo-HTT^Q23^-associated proteins (**Fig. S5J**). This pattern is notably also pronounced among the high-confidence clusters from the IP-MS groups (**Fig. 3M and N, Fig. S5K and L**). Together these findings suggest that HTT-HAP40 heterooligomers through PPIs function as stabilizers for associated proteins involved in rRNA processing, translation and the assembly of cytoskeletal structures and loss of HTT or HAP40 function significantly decreases their cellular abundance.

### Loss of HAP40 leads to widespread transcriptional changes and upregulation of multiple CLEAR pathway-associated proteins in striatal cells

We next investigated the effects of HAP40 loss on gene expression programs in striatal cell models that endogenously produce either wild-type or mutant full-length HTT. Principal component analysis of transcriptome data sets revealed divergent gene expression profiles in striatal cell lines between ST*Hdh*^Q7^ and ST*Hdh*^Q111^ cells, with further divergence observed upon HAP40 loss in each background (**Fig. 4A**). Strikingly, differential gene expression analysis revealed >11,000 significantly changed transcripts (adj p < 0.05) when data sets of ST*Hdh*^Q7^ and ST*Hdh*^Q7^-*Hap40*KO cell lines were compared (**Fig. 4B, Table S3**), indicating that HAP40 loss exerts a profound impact on the transcriptome. In ST*Hdh*^Q7^-*Hap40*KO cells, the abundance of ∼5,600 genes were significantly increased or decreased relative to the parental ST*Hdh*^Q7^ line (**Fig. 4B**), demonstrating both activation and repression of transcriptional programs. Notably, the number of differentially expressed genes (DEGs) was significantly higher in the mutant background, implying that cells producing mutant HTT^Q111^ may experience greater transcriptional stress upon HAP40 depletion than those expressing HTT^Q7^. IPA of DEGs of ST*Hdh*^Q7^ and ST*Hdh*^Q7^-*Hap40*KO cells revealed dysregulation across a wide range of pathways and cellular processes (**Fig. 4C**). Among the top 10 significantly upregulated pathways in ST*Hdh*^Q7^-*Hap40*KO and ST*Hdh*^Q111^-*Hap40*KO (**Fig. 4C and Fig. S7A)** cells, we identified the CLEAR (Coordinated Lysosomal Expression and Regulation) gene regulatory network (**Fig. 4D**). This transcriptional program, which is induced by overproduction of the transcription factor TFEB(37), governs autophagy and lysosomal biogenesis(38, 39), suggesting that HAP40 functions as a proteostasis maintenance regulator and its loss activates protein degradation pathways in striatal cells. Closer inspection of CLEAR network-associated genes confirmed the upregulation of genes encoding key lysosomal proteins (**Fig. 4E).**

**Fig. 4.**
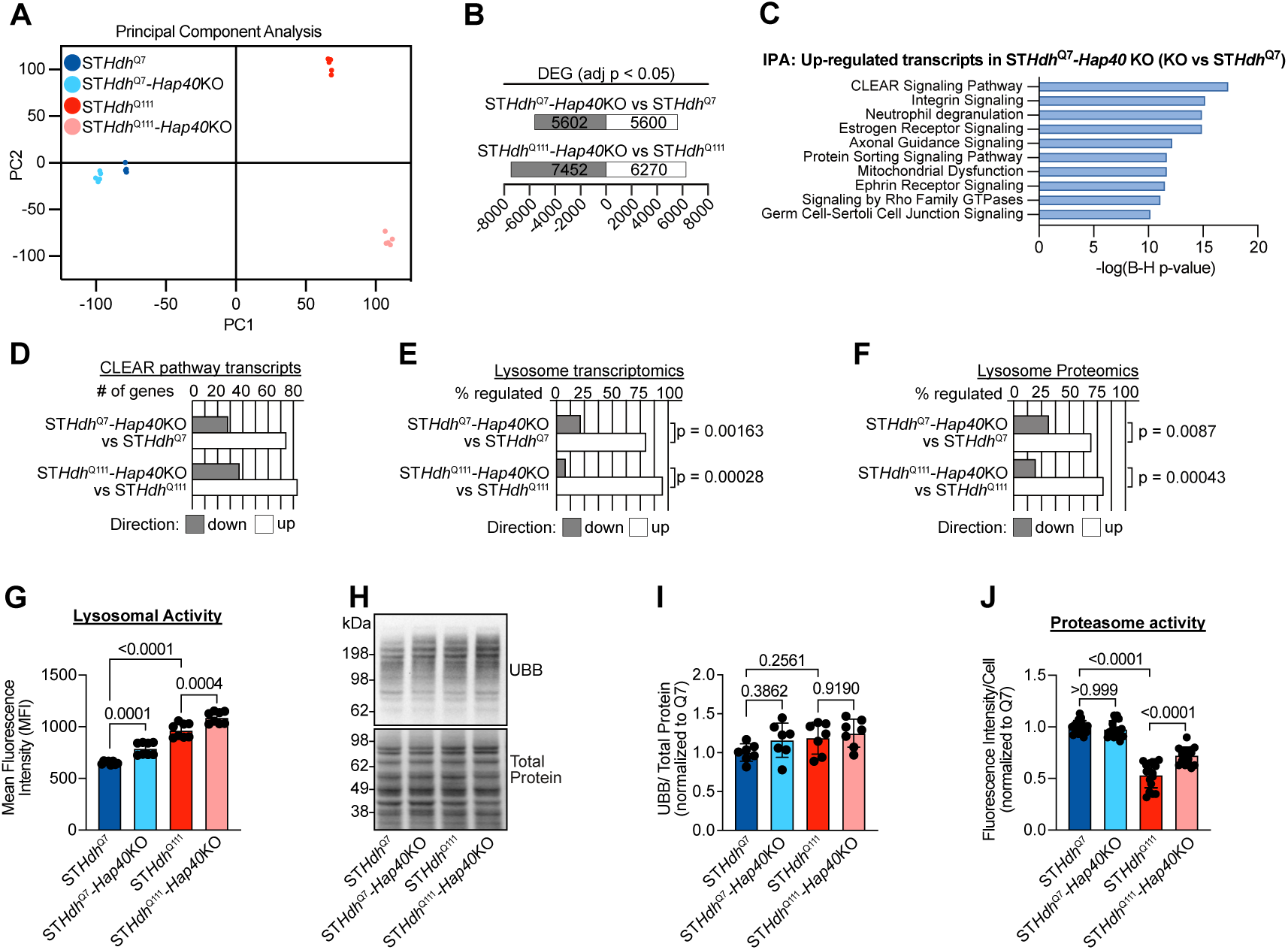
Transcriptional and lysosomal remodeling in wild-type and mutant HTT striatal cells lacking HAP40. **A)** Principal component analysis (PCA) of transcriptome profiles from ST*Hdh*^Q7^, ST*Hdh*^Q7^-*Hap40*KO, ST*Hdh*^Q111^, and ST*Hdh*^Q111^-*Hap40*KO cell lines. Each point represents a biological replicate (n = 4) and is colored by genotype; the plot visualizes global transcriptional differences among the conditions. **B)** Numbers of significantly upregulated (white) and downregulated (grey) transcripts (adjusted p < 0.05) in the two *Hap40*KO lines relative to their respective parental controls. The total number of differentially expressed genes is shown in each bar. **C)** Ingenuity Pathway Analysis (IPA) of transcripts significantly upregulated (adjusted p ≤ 0.05) in ST*Hdh*^Q7^-*Hap40*KO versus ST*Hdh*^Q7^ cells. Canonical pathways are ranked by Benjamini–Hochberg FDR-adjusted p-values. **D)** Panel shows the number of genes that belong to the CLEAR (Coordinated Lysosomal Expression and Regulation) network and are either increased or decreased in the two different striatal knockout cell lines compared with their respective parental (wild-type) lines. For each genotype, ST*Hdh*^Q7^-*Hap40*KO and ST*Hdh*^Q111^-*Hap40*KO, we identified transcripts whose expression changed significantly (adjusted p ≤ 0.05). Those significant transcripts were then matched against a curated list of CLEAR-pathway genes taken from the IPA database. The resulting counts of upregulated and downregulated CLEAR genes are presented side-by-side for each knockout condition. **E)** Percentage of lysosomal genes that are significantly up- or downregulated in the two *Hap40*KO lines relative to controls. For each KO line, we first identified all lysosomal genes that passed the significance threshold and then expressed the result as a percentage of the detected lysosomal genes in the dataset. Upregulation of lysosomal genes is significantly overrepresented relative to downregulation, in contrast to the up/down-distribution observed in the total dataset (one-sided Fisher’s exact test). **F)** Same analysis as in **E**, but for lysosomal proteins quantified by proteomics. **G)** Lysosomal activity was measured by flow cytometry, and mean fluorescence intensity (MFI) was recorded. Data are presented as mean ± SD of two independent experiments with four technical replicates of 10,000 cells each. Statistical significance was evaluated using one-way ANOVA with Šídák’s multiple comparisons test. **H)** Representative immunoblot of mouse striatal total lysates probed for polyubiquitin and no-stain labelling as protein loading control. **I)** Quantification of polyubiquitin levels relative to total protein from no-stain labelling. Data were normalized to ST*Hdh*^Q7^. Data are presented as mean ± SD of two independent experiments with 3-5 technical replicates each. Statistical significance was assessed by one-way ANOVA with Tukey’s multiple comparison test. **J)** Quantification of proteasome activity assay performed with mouse striatal cell lines. Data are presented as mean ± SD of four independent experiments with four technical replicates each. Statistical significance was assessed by one-way ANOVA with Bonferroni’s multiple comparison test.

Furthermore, we performed a total proteome analysis on the striatal cell lines (**Fig. S7B-D** and **Table S4**) and observed increased levels of lysosomal proteins in ST*Hap40*KO cells in comparison to controls (**Fig. 4F**). Thus, our analysis of both transcriptome and proteome data sets indicates that *Hap40*KO in striatal cells activates the autophagy-lysosome pathway(40), a major subcellular process responsible for degrading long-lived proteins(41) or dysfunctional organelles.

Next, we assessed the lysosomal activity in striatal protein extracts by quantifying the proteolytic cleavage of a lysosome-specific self-quenched substrate. We found that substrate cleavage is significantly higher in ST*Hdh*^Q7^-*Hap40*KO and ST*Hdh*^Q111^-*Hap40*KO cell lines compared to controls (**Fig. 4G**), suggesting that transcriptional activation of CLEAR genes (**Fig. 4D**) enhances lysosomal protein degradation in striatal cells. The abundance of ubiquitinated proteins in cell extracts prepared from striatal cell lines was not significantly altered by HAP40 loss (**Fig. 4H and I**). However, a slight increase in proteasome-activity was observed in ST*Hdh*^Q111^-*Hap40*KO cells compared to ST*Hdh*^Q111^ cells (**Fig. 4J**), indicating that loss of HAP40 weakly activates the ubiquitin proteasome pathway in ST*Hdh*^Q111^-*Hap40*KO cells. Together these results indicate that HAP40 functions as a proteostasis regulator in striatal cells and controls transcriptional programs that regulate protein degradation pathways.

### Loss of HAP40 leads to an accumulation of the autophagy-associated proteins p62, LC3B-II and STX17 in ST*Hdh*^Q111^-*Hap40*KO cells

We explored whether changes in lysosomal activity (**Fig. 4G**) may influence the abundance of the autophagy receptor p62(42), whose steady state levels in cells are controlled by the activity of the autophagy-lysosomal protein degradation pathway(43). Analysis of protein extracts by SDS-PAGE and immunoblotting revealed that steady state levels of p62 are slightly lower in ST*Hdh*^Q7^-*Hap40*KO than in ST*Hdh*^Q7^ cells (**Fig. 5A and B**), suggesting that CLEAR pathway activation (**Fig. 4D**) reduces p62 abundance and increases its degradation in striatal *Hap40*KO cells. Interestingly, significantly higher p62 levels were quantified in ST*Hdh*^Q7^ than in ST*Hdh*^Q111^ cells, indicating that endogenous production of HTT^Q111^ increases lysosome activity (**Fig. 4G**) and enhances p62 degradation through the autophagy-lysosomal protein degradation pathway (**Fig. 5B**). However, to our surprise we observed that steady state levels of p62 in ST*Hdh*^Q111^-*Hap40*KO cells in comparison to ST*Hdh*^Q111^ cells were significantly increased (**Fig. 5A and B**), indicating that HAP40 loss in cells with pathogenic HTT^Q111^ impairs selective autophagy and leads to accumulation of p62, although lysosome activity is high (**Fig. 4G**).

**Fig. 5.**
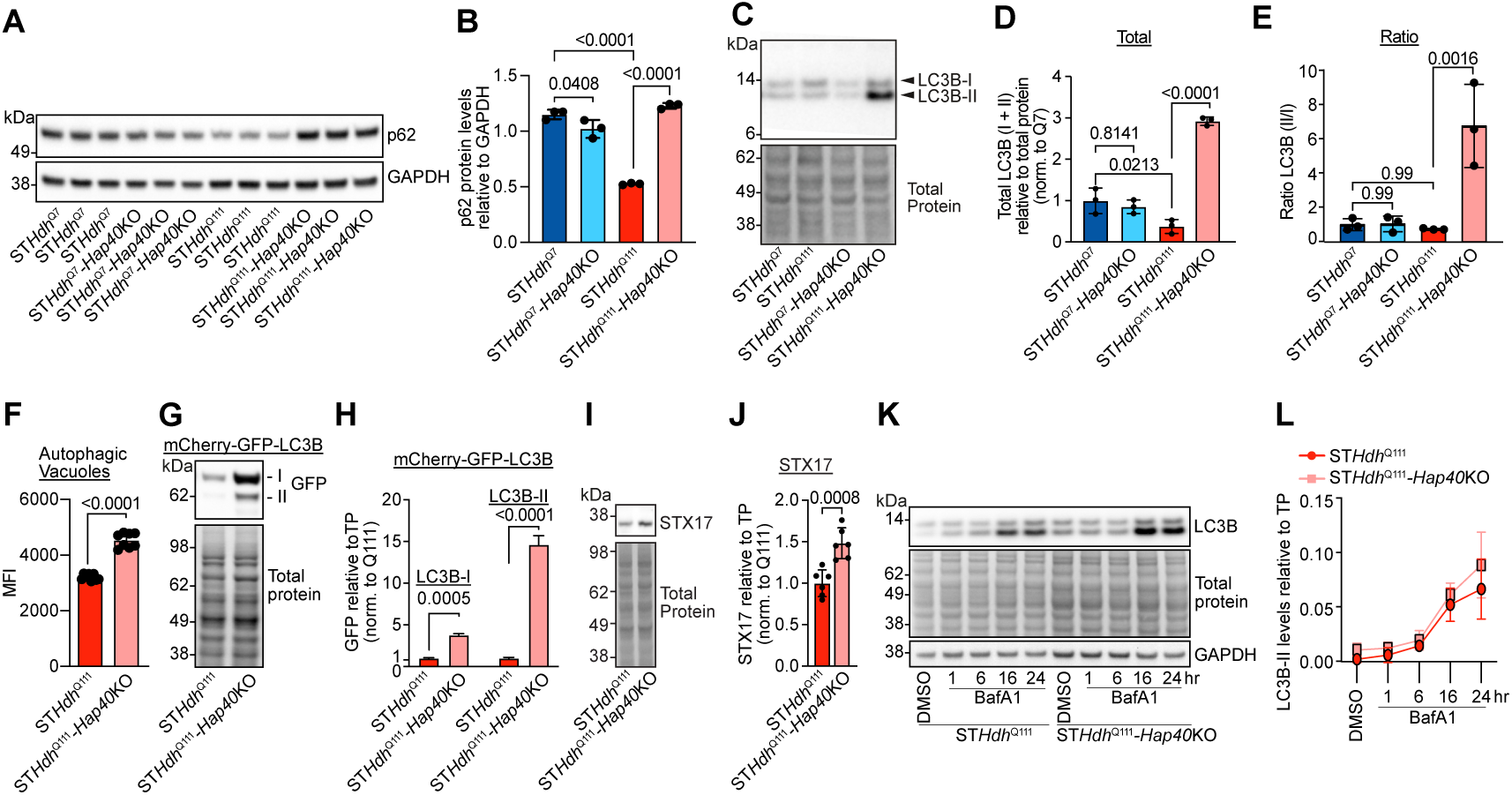
Loss of HAP40 results in impaired autophagosome-lysosome flux. **A**) Representative immunoblots of different mouse striatal total cell lysates. Membranes were probed with antibodies against p62 and the loading control GAPDH. **B)** Quantification of p62 protein levels in **A**, expressed relative to GAPDH. Data are presented as mean ± SD of four biological replicates. Statistical significance was determined by one-way ANOVA with Tukey’s multiple comparisons test. **C)** Representative immunoblot of mouse striatal lysates probed for LC3B with total protein staining as loading control. **D)** Quantification of total LC3B I & II protein bands relative to total protein and **E)** Quantification of ratio of LC3BII/LC3BI, both related to **C**. For both **D** and **E** data are presented as mean ± SD of three biological replicates. Statistical significance was determined by one-way ANOVA with Tukey’s multiple comparisons test. **F)** Quantification of autophagic vacuoles was performed using an autophagy assay kit and measured by flow cytometry. Data are presented as the mean fluorescence intensity (MFI) ± SD of two independent experiments, each with four technical replicates of 10,000 cells. Statistical significance was assessed by one-way ANOVA with Tukey’s multiple comparisons test. **G)** Representative immunoblot of mouse striatal lysates transfected with pDEST-mCherry-GFP-LC3B and probed using an anti-GFP antibody, with total protein staining as loading control. **H)** Quantification of total mCherry-GFP-LC3B-I and -II protein bands relative to total protein (TP). Data are presented as mean ± SD of three biological replicates. Statistical significance was assessed by Welch’s t-test. **I)** Representative immunoblot of striatal lysates probed with anti-STX17, and total protein staining. **J)** Quantification of STX17 relative to total protein (TP); data are presented as mean ± SD of three biological replicates. Statistical significance was assessed by Welch’s t-test. **K)** Representative immunoblot of mouse striatal lysates treated with 100 mM Bafilomycin A1 (BafA1) or an equal volume of DMSO for various time points. **L)** Measurement of LC3B-II levels relative to total protein (TP) and across BafA1 treatment timepoints (n=2).

To assess whether loss of HAP40 in ST*Hdh*^Q111^-*Hap40*KO cells alters autophagic flux(44), we next quantified the lipidated marker protein LC3B-II(45), which measures the abundance of autophagic vesicles in neuronal cells(46). Analysis of protein extracts by SDS-PAGE and immunoblotting revealed significantly higher LC3B-II levels and a higher LC3B-II/I ratio in ST*Hdh*^Q111^-*Hap40*KO than in ST*Hdh*^Q7^, ST*Hdh*^Q111^ and ST*Hdh*^Q7^-*Hap40*KO cells (**Fig. 5C-E**), indicating that autophagic flux is impaired and autophagic vesicles accumulate in HTT^Q111^-producing striatal cells when HAP40 is absent. This was also confirmed by immunofluorescence microscopy studies showing a significantly higher number of LC3B-positive puncta in ST*Hdh*^Q111^-*Hap40*KO than in ST*Hdh*^Q111^ control cells (**Fig. S8A and B**). Furthermore, treatment of cells with a fluorescent dye that selectively labels autophagic vacuoles revealed higher signals in ST*Hdh*^Q111^-*Hap40*KO than in ST*Hdh*^Q111^ cells (**Fig. 5F**), supporting the hypothesis that autophagic vesicles accumulate in HTT^Q111^-producing *Hap40*KO cells. To quantify autophagic LC3B degradation, we also expressed a tandem monomeric mCherry-GFP-tagged LC3B(47) fusion protein in striatal cells. Notably, higher amounts of mCherry-GFP-LC3B (both I and II forms) were detectable in ST*Hdh*^Q111^-*Hap40*KO cells compared to ST*Hdh*^Q111^ control cells (**Fig. 5G and H**), confirming that autophagic protein degradation is decreased in the absence of HAP40. A key step in the delivery of cargo to the lysosome is the fusion of autophagosomes with lysosomes. This step gets facilitated by STX17, an autophagosome SNARE protein, which is recruited to mature autophagosomes^50,48^(48). Immunoblotting revealed increased STX17 levels in ST*Hdh*^Q111^-*Hap40*KO cells compared to ST*Hdh*^Q111^ cells (**Fig. 5I and J**), indicating that autophagosome-lysosome fusion is impaired in HTT^Q111^-producing cells, when HAP40 is absent.

Finally, investigations with Bafilomycin A1 (BafA1), a specific inhibitor of V-type H+-ATPase(49) that potently blocks autophagosome-lysosome fusion(50) were performed to assess the impact of HAP40 loss on the rate of autophagosome formation in striatal cells. Treatment with BafA1 revealed similar kinetics of LC3B-II accumulation in both mutant HTT cell lines (**Fig. 5K and L**), demonstrating that loss of HAP40 in HTT^Q111^-producing striatal cells does not significantly influence the rate of autophagosome biogenesis. Together, these studies indicate that HAP40 function is critical for the autophagosome-lysosome fusion and loss of this function in ST*Hdh*^Q111^-*Hap40*KO cells leads to an impairment of autophagic flux and an abnormal accumulation of autophagic vesicles.

### Loss of HAP40 promotes the release of pathogenic HT**T^Q11^**^1^ into the extracellular space

Previous studies indicate that full-length HTT is an autophagy substrate that gets readily degraded in mammalian cells by the autophagosome-lysosome pathway(51, 52). Therefore, we next investigated the impact of HAP40 depletion on the abundance of HTT in striatal cell lines. Striatal cell lines were first treated with BafA1 to assess HTT accumulation when autophagosome-lysosome fusion is fully blocked(45). As expected, we observed a significant increase of HTT^Q7^, when ST*Hdh*^Q7^ or ST*Hdh*^Q7^-*Hap40*KO cells were treated with BafA1 (**Fig. 6A and B**), confirming previous observations that the autophagy-lysosome pathway degrades full-length HTT(53). However, surprisingly no significant accumulation of HTT^Q111^ was detectable in BafA1 treated ST*Hdh*^Q111^ and ST*Hdh*^Q111^-*Hap40*KO cells (**Fig. 6A and B**), indicating that this protein is not cleared by canonical autophagy(54). Thus, alternative pathways such as the previously described non-canonical secretory autophagy pathway(55), which maintains proteostasis upon lysosome inhibition, might be activated in HTT^Q111^-producing striatal cells.

**Fig. 6.**
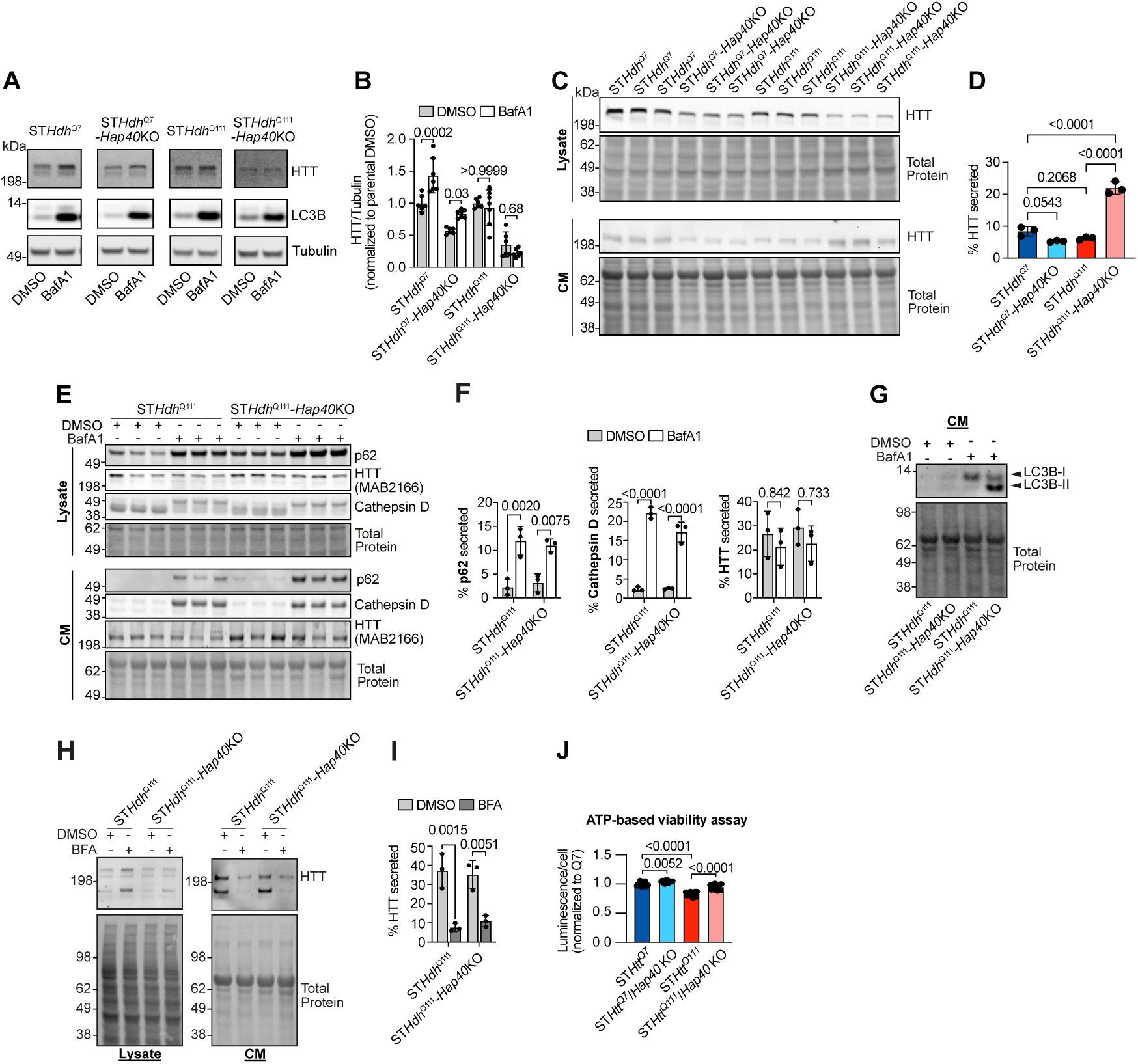
Loss of HAP40 stimulates mHTT secretion in ST*Hdh*^Q111^-*Hap40*KO striatal cells. **A)** Representative immunoblots of total cell lysate treated with DMSO or 100 mM BafA1 for 24 hours. **B)** Quantification of HTT protein levels relative to tubulin, related to A. Data were normalized to parental control (either ST*Hdh*^Q7^ or ST*Hdh*^Q111^). Data are presented as mean ± SD of two independent replicates, each with three biological replicates. Statistical significance was assessed by two-way ANOVA with Bonferroni’s multiple-comparisons test. **C)** Representative immunoblots of striatal lysates and conditioned media (CM), probed with anti-HTT (MAB2166) and total protein staining. **D)** Quantification assessing the percentage (%) of secreted protein in conditioned media. Data are presented as mean ± SD of three biological replicates. Statistical significance was assessed by one-way ANOVA with Tukey’s multiple comparisons test**. E)** Immunoblot of striatal lysates and conditioned media (CM) in the presence of 100 mM BafA1 or DMSO for 24 hours, probed with anti-p62, anti-HTT (MAB2166), anti-Cathepsin D, and total protein staining. **F)** Quantification assessing the percentage (%) of secreted protein in conditioned media. Data are presented as mean ± SD of three biological replicates. Statistical significance was assessed by two-way ANOVA with Tukey’s multiple comparisons test**. G)** Representative immunoblot of striatal conditioned media treated for 24 hours with 100 mM BafA1 or equal volume of DMSO, probed with anti-LC3B and total protein staining. **H)** Representative immunoblots of striatal lysates and conditioned media (CM) in the presence of 5 µg/mL Brefeldin A (BFA) or DMSO for 4 hours, probed with anti-HTT (MAB2166) and total protein staining. **I)** Quantification assessing the percentage (%) of HTT secreted protein in conditioned media. Data are presented as mean ± SD of three biological replicates. Statistical significance was assessed by two-way ANOVA with Tukey’s multiple comparisons test**. J)** Quantification of ATP-based viability assay based on the luminescence signal relative to the number of cells per well, normalized to ST*Hdh*^Q7^. Data represent the mean ± SD of two independent experiments, each with 7-8 technical replicates. Statistical significance was assessed by two-way ANOVA with Bonferroni’s multiple-comparisons test.

To address this question, we next prepared crude protein fractions from striatal cells and conditioned media and analyzed them by SDS-PAGE and immunoblotting using the anti-HTT MAB2166 antibody. As expected, we observed that HTT^Q7^ protein levels in lysates prepared from ST*Hdh*^Q7^-*Hap40*KO cells are significantly lower than in ST*Hdh*^Q7^ cells (**Fig. 6C**), confirming our initial observations that loss of HAP40 significantly decreases intracellular HTT protein levels (**Fig. 1C**). A similar result was obtained when cell extracts of ST*Hdh*^Q111^ cells and ST*Hdh*^Q111^-*Hap40*KO cells were analyzed, although overall HTT^Q111^ protein levels were significantly lower than HTT^Q7^ protein levels in striatal cells. Strikingly, when conditioned medium was analyzed, we detected significantly higher amounts of HTT^Q111^ in fractions in ST*Hdh*^Q111^-*Hap40*KO cells compared to the other investigated cell lines (**Fig. 6C and 6D**), indicating that the pathogenic protein in the absence of HAP40 indeed gets secreted more efficiently than in its presence. Importantly, low levels of HTT^Q7^ were also released in ST*Hdh*^Q7^ cells, demonstrating that both mutant and wild-type HTT are secreted. However, loss of HAP40 in these cells did not exaggerate HTT^Q7^ secretion, suggesting that most wild-type protein likely gets degraded intracellularly potentially through the autophagy-lysosomal pathway (**Fig. 6A and 6B**). Cumulatively, these studies demonstrate that pathogenic HTT^Q111^ neither in the presence of HAP40 nor in its absence is degraded by canonical autophagy in striatal cells. However, intracellular stress (**Fig. 4A-D**) induced by HAP40 loss enhances HTT^Q111^ secretion into the extracellular space and reduces its intracellular abundance.

To determine the secretory pathway(56) responsible for secretion of HTT^Q111^ in striatal cells, we employed different compounds that inhibit critical proteins involved in secretion. We first treated cells with BafA1, a potent inhibitor of lysosome activity that stimulates secretory autophagy. Interestingly, we observed that BafA1 treatment neither influences HTT^Q111^ secretion in ST*Hdh*^Q111^-*Hap40*KO nor in ST*Hdh*^Q111^ cells (**Fig. 6E and F**), indicating that lysosome inhibition-induced secretory autophagy does not influence the release of HTT^Q111^ into the extracellular space. However, BafA1 treatment significantly increased the secretion of p62, cathepsin D and LC3-II into the conditioned medium of striatal cell lines (**Fig. 6E-G**), confirming previous observations that lysosome inhibition stimulates secretory autophagy(55).

Next, we investigated whether trafficking through the ER-to-Golgi route influences HTT^Q111^ secretion. Striatal cells were treated with Brefeldin A (BFA), an ER-to-Golgi protein trafficking inhibitor that impairs conventional secretion of proteins with N-terminal signal sequences but also can inhibit secretion of proteins without such sequences(57, 58). Treatment with BFA resulted in intracellular accumulation of HTT^Q111^ and potently decreased the abundance of the mutant protein in the conditioned medium (**Fig. 6H and I**), suggesting that molecular events that facilitate canonical ER-to-Golgi trafficking are critical for the release of mHTT into the extracellular space.

Lastly, we quantified ATP levels to assess whether increased secretion reduces mutant HTT^Q111^ toxicity in ST*Hdh*^Q111^-*Hap40*KO striatal cells. We found that ATP levels were significantly increased in ST*Hdh*^Q111^-*Hap40*KO in comparison to control ST*Hdh*^Q111^ cells (**Fig. 6J**), indicating that decreasing intracellular HTT^Q111^ protein levels through secretion reduces mutant HTT-induced cellular stress and improves protein homeostasis.

## Discussion

In this study we investigated the function of the α-helical, tetratricopeptide repeat protein HAP40 in mammalian cells as well as its influence on wild-type HTT function and the pathobiology of mHTT. HAP40 is a highly conserved, direct binding partner and potential key regulator of mHTT toxicity and therefore a highly relevant possible target in HD therapy. Utilizing imaging and biochemical methods, we found that HAP40 is an obligate HTT interaction partner under endogenous conditions (**Fig. 1G-K),** meaning that it is permanently associated with HTT. Our studies suggest that HAP40’s biological role in cells is to control HTT function through its binding.

One aspect of HTT is that it is reported to bind thousands of proteins(59, 60), a unique feature that was attributed to the ability of HTT to operate in different pathways like a Swiss-army knife. To date, there are no interactions studies that have assessed the impact of HAP40 on HTT interactions. Our co-immunoprecipitation experiments followed by mass spectrometry revealed that apo-HTT in the absence of HAP40 binds more cellular proteins than HTT-HAP40 heterooligomers (**Fig. 3C**), supporting the hypothesis that HAP40 binding to HTT controls its association with other cellular proteins. We suggest that HAP40 regulates HTT interactions by stabilizing its 3D conformation and defining its cellular localization. While HAP40 binding has been reported previously to stabilize the structure of full-length HTT(16, 61), we show for the first time, utilizing the sensor protein NL-HTT^Q23(2689-mCit)^ and quantitative *in-cell* BRET measurements, that HAP40 stabilizes full-length HTT conformation in cells. We found that the molecular distance between N-and C-termini in HTT of stable HTT-HAP40 heterooligomers is shorter than in full-length apo-HTT molecules, which may expose unique interaction surfaces due to its altered structure (**Fig. 2**). Additionally, loss of HAP40 altered the subcellular distribution of full-length HTT. While HTT-HAP40 complexes in HEK293 cells were concentrated in the perinuclear region, the distribution of apo-HTT was more dispersed (**Fig. S2K**). This suggests that it has the potential to interact with many other cellular proteins in the absence of HAP40. We propose that HAP40 binding enables HTT to localize to specific subcellular locations and to expose interaction surfaces that are critical for its molecular function and specific association with other cellular proteins.

Our GO term enrichment analysis of interaction data sets supports the hypothesis that apo-HTT and HTT-HAP40 heterooligomers bind distinct proteins in mammalian cells. While proteins involved in transmembrane transport processes, lipid biosynthesis or mitochondrial functions (**Fig. 3D**) were found to predominantly associate with apo-HTT, proteins involved in ribosome biogenesis and rRNA metabolism were preferentially co-enriched with HTT-HAP40 heterooligomers. We propose that multiprotein complexes with HTT-HAP40 heterooligomers fulfill specific functional tasks in cells, while protein complexes with apo-HTT are potentially deleterious for cells, because they are non-physiological or unregulated interactions. Hence, the association of apo-HTT with proteins implicated in vesicle transport processes or mitochondrial functions may contribute to cellular dysfunction and stress, while the interactions of HTT-HAP40 heterooligomers with proteins involved in translation and/or ribosome biogenesis likely promote these specific subcellular processes. An association of HTT with proteins in these subcellular processes has also has been reported previously (62, 63), supporting our observations with HTT-HAP40 heterooligomers. Functional investigations in cell models(64) have shown that full-length HTT directly binds RNA(65) and facilitates protein synthesis(62, 64).

Our proteomics studies further support the notion of HAP40 as a HTT PPI regulator. The comparison of HTT interaction and proteomics data sets revealed that a large fraction of the identified HTT-HAP40-associated proteins is significantly decreased in abundance in HEK293-HAP40KO and HEK293-HTTKO cells (**Fig. 3K–3N**). This indicates that HTT-HAP40 complexes stabilize interacting proteins under physiological conditions and HAP40 or HTT loss leads to decreased steady-state levels. A prime example is the neurofilament medium (NEFM) protein. This protein is enriched with the HTT-HAP40 heterooligomers in IP experiments, but upon loss of HTT or HAP40, its protein abundance is decreased. A previous proteomic study in an HD mouse model identified NEFM exhibiting a significant correlation with HTT abundance, supporting our finding(60). This suggests that HTT-HAP40 heterooligomers are required to stabilize associated interactors and protect them from being degraded.

Analysis of gene regulatory networks and biochemical studies position HAP40 as proteostasis regulator. The transcriptome profiles of HAP40 KO and control striatal lines revealed a massive transcriptional dysregulation of thousands of genes leading to reprogramming of multiple cellular pathways upon the loss of HAP40 (**Fig. 4B**). We observed that the transcriptional dysregulation in HTT^Q111^-producing HAP40 KO cells was significantly more pronounced than in HTT^Q7^-producing HAP40 KO cells (**Fig. 4B**), indicating that, in the absence of HAP40, mHTT is more proteotoxic for cells and induces a stronger transcriptional response than the respective wild-type protein. Intriguingly, our analysis of gene expression data revealed an activation of the CLEAR (Coordinated Lysosomal Expression and Regulation) network (**Fig. 4C and S7A**) in HAP40 KO cells, which regulates the activity of autophagic protein degradation through the lysosomal pathway in mammalian cells(37). Therefore, HAP40 may function as proteostasis regulator that controls the activity of protein degradation pathways under physiological conditions. When this function is lost, compensatory cellular programs are activated to re-adjust protein homeostasis. This view is also supported by our functional studies with HAP40 KO and control cell lines, demonstrating that loss of HAP40 significantly increases lysosome activity in striatal cells (**Fig. 4G**). In addition, we observed an abnormal accumulation of the autophagy marker proteins p62, LC3-II and STX17 in ST*Hdh*^Q111^-*Hap40*KO cells (**Fig. 5A-J**), which is an indication of autophagic flux impairment. Importantly, previous investigations suggest that HTT plays a functional role in selective autophagy(66, 67), which is in agreement with our observations that loss of HAP40 in HTT^Q111^-producing striatal cells impairs autophagy. Also, an impairment of autophagy has been shown iPSC-derived neurons of HD patients endogenously producing a pathogenic HTT protein(4). Our studies also underline reports that full-length HTT acts as a regulator of gene expression(68, 69) and influences mRNA splicing(6). In summary, we propose that HAP40 serves as a proteostasis regulator by controlling of HTT interactions and gene regulatory networks. In this function it maintains the steady-state levels of thousands of cellular proteins and governs the activity of lysosomal genes (**Fig. 7**).

**Fig. 7.**
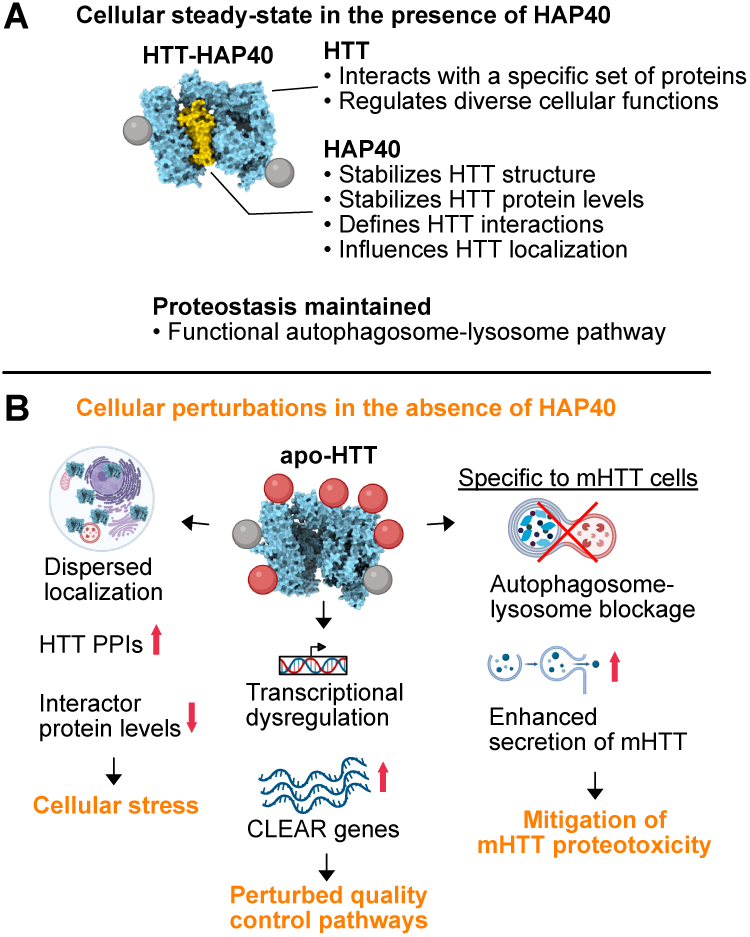
Study model of HAP40 function. **A)** The HTT-HAP40 heterooligomer governs the stability and functional repertoire of HTT. In the presence of HAP40, HTT adopts a stable conformation, localizes to specific subcellular structures, controls the level of associated proteins, and defines which proteins interact with HTT. A primary outcome of HAP40 regulation of HTT is a preservation of autophagosome-lysosome flux and a control of lysosomal activity. **B)** In contrast, loss of HAP40 alters HTT localization and destabilizes apo-HTT conformation, leading to unregulated protein-protein interactions and altered interactor protein levels leading to cellular stress. Additionally, loss of HAP40 causes transcriptional dysfunction, resulting in the upregulation of CLEAR network genes composed primarily of lysosomal genes, that control the activity of quality-control pathways. In the presence of mHTT, loss of HAP40 leads to impaired autophagosome-lysosome flux, to which cells respond by enhancing secretion to mitigate mHTT proteotoxicity. Created in BioRender. Wanker, E. (2026) https://BioRender.com/smle33q.

The activation of lysosome-based protein clearance pathways (**Fig. 4G**) may explain our observations that the abundance of full-length HTT is significantly lower in cells without HAP40 than in controls (**Fig. 1A-D**). Surprisingly however, our experimentation showed that the autophagic flux in ST*Hdh*^Q111^-*Hap40*KO cells is impaired and that mutant HTT^Q111^ likely is not cleared through this degradation pathway. As previous investigations showed that cells can secrete mHTT(70, 71), we hypothesized that striatal ST*Hdh*^Q111^-*Hap40*KO cells might decrease intracellular HTT^Q111^ protein levels through activation of secretory pathways. Strikingly, our analysis of conditioned media showed that ST*Hdh*^Q111^-*Hap40*KO cells secrete significantly higher amounts of HTT^Q111^ than ST*Hdh*^Q111^ cells (**Fig. 6C and D**), supporting the idea that secretory pathways are indeed activated in HAP40 KO cells. We conclude that upon impairment of intracellular protein degradation pathways proteotoxic proteins such as HTT^Q111^ get secreted more efficiently into the extracellular space to maintain protein homeostasis.

Interestingly, the release of mHTT into the extracellular space was blocked when striatal cells were treated with the secretion inhibitor brefeldin A (BFA) (**Fig. 6H and I**). BFA inhibits ARF1 activation at ER and Golgi membranes(72) and disrupts protein trafficking from the ER to the Golgi(73), suggesting that secretion through the Golgi apparatus is critical for the release of mHTT into the extracellular space. This is supported by previous reports that HTT preferentially associates with ER and Golgi membranes through its conserved N-terminal N17 domain(23). Furthermore, HTT function was shown to be required for efficient ER-to-Golgi transport and the fusion of secretory vesicles with the plasma membrane(74). Thus, our results suggest that mHTT, which directly binds to membranes as well as interacts with various Golgi-associated proteins(75, 76), might be released into the extracellular space by an unconventional secretory route similar to the Golgi-associated protein GRASP55(77). Interestingly, GRASP55 loss was shown to effect the secretion of a short mHTT fragment (78). More detailed investigations are necessary to elucidate the specific molecular events that facilitate secretion of full-length mHTT.

We propose that our observations with striatal cell lines have important implications for a better understanding of the disease mechanism in HD and the development of future therapeutic strategies. Our finding that HAP40 functions as a proteostasis regulator in cells, determining the steady-state levels of thousands of cellular proteins, suggests that therapeutic strategies targeting HAP40 for protein lowering, such as siRNAs or ASOs(79), may not be a beneficial strategy. Although, lowering HAP40 could lead to a decrease in mHTT levels, it poses the risk of also reducing the levels of many other cellular proteins. Moreover, decreasing HAP40 protein levels with therapeutic molecules might activate comprehensive transcriptional programs and stress pathways to re-adjust protein homeostasis. Additionally, knock-down of HAP40 in neurons could also promote the secretion of mHTT into the extracellular space, which in turn might stimulate mHTT spreading, a phenomenon that has been observed in *Drosophila*(80) and cell models(81). Spreading of misfolded disease proteins such as tau or α-synuclein is associated with dysfunction and neurotoxicity in neurodegenerative diseases, including Alzheimer’s disease (AD), Parkinson’s disease (PD), and amyotrophic lateral sclerosis (ALS)(82, 83). Taken together, our findings define HAP40 as a proteostasis regulator that is required to stabilize canonical HTT interactions and to mitigate mutant HTT proteotoxicity.

## Materials and Methods

### Cell lines

Human embryonic kidney line 293 (HEK293) wild-type and HEK293-HTTKO cells(84) were grown in Dulbecco’s modified Eagle’s medium (Thermo Fisher Scientific, 41965) supplemented with 10% heat-inactivated fetal bovine serum (Thermo Fisher Scientific, 10500064), and 1% penicillin and streptomycin (Thermo Fisher Scientific, 15140122) at 37°C, and 5% CO_2_. Mouse striatal (ST) *Hdh*^Q7^ (ST*Hdh*^Q7/Q7^, CH00097) and ST*Hdh*^Q111^ (ST*Hdh*^Q111/Q111^, CH00095) neuronal cell lines(28) were obtained from the HD community Biorepository at Coriell Institute. Striatal cells were maintained in complete DMEM medium (Thermo Fisher Scientific, 41965) supplemented with 10% heat-inactivated fetal bovine serum, 1% penicillin/streptomycin, 0.4 mg/mL geneticin (Thermo Fisher Scientific, 10131035), 1x MEM non-essential amino acids (Thermo Fisher Scientific, 11140035) and incubated at 33°C and 5% CO_2_. After thawing, cells were cultured for one week to reach log phase before cell pellets were collected or experiments were performed. Cells were subcultured every three to four days.

### CRISPR-Cas9 engineering

The HEK293-HAP40KO-clone1B4 (HEK293-HAP40KO) cell line was generated using CRISPR-Cas9 genome editing to induce indels close to the start triplet ‘ATG’ of the human *HAP40* (*F8A1*) gene. A single gRNA (sgRNA) sequence (ACCCGAGGCCGGGGACTTCC) was cloned into pSpCas9(BB)-2A-Puro (addgene: 62988) and was used for gene editing and selection of cells expressing puromycin cassette (Puro). One million cells were seeded into a six-well plate and 2 µg of plasmid was transfected into cells using FuGENE transfection reagent (Promega, E2311) at a 3:1 reagent µL:µg DNA ratio. 24 hours later, cells were selected for uptake of Cas9 plasmid by adding puromycin (1 µg/mL final). After 3 days of puromycin selection, single-cells were obtained by serial dilution into 96-well plates, one cell per every three wells. Single-cell colonies were expanded and PCR amplification was performed targeting the edited site using the following primers, forward primer: CTTTAGCAACCTAGACCAC; reverse primer: CACATGAGGAGTACA-AGAGTAG. The expected PCR product was purified and subjected to Sanger sequencing with the same PCR oligos. Sequencing confirmed that the clonal cell line had a frameshift mutation resulting in a premature stop codon. Immunoblot analysis using 25 µg of total cell lysate and an anti-HAP40 antibody (Atlas, HPA046960,1:500) confirmed the loss of full-length HAP40 expression in KO clones.

HEK293 HTT-mNeonGreen-clone1B (HEK293-HTT-mNG) knock-in cell line was generated using a similar approach with CRISPR-Cas9 editing. In this case, a single guide RNA (sgRNA) targeting the 3’ end of the human *HTT* gene (CCACCTGCTGAGCGCCATGG) was co-transfected with a custom-designed homology-directed repair template. This template contained a human codon-optimized mNeonGreen gene flanked by 500 base pairs of the HTT insertion site. Cells that exhibited mNeonGreen fluorescence were collected via fluorescence-activated cell sorting and serially diluted to obtain single-cell colonies. These colonies were subjected to PCR amplification using the following primers: GCTGGAGCAGGTGGACGTGAAC and TCTGGAA-GGCCTCAGGCTCAGC. An unedited clone produced an amplified DNA fragment of approximately 416 base pairs, while an edited knock-in clone amplified a 1154 base pair fragment. Clones displaying only the 1154 base pair band (indicating homozygous knock-in) were further analyzed by Sanger sequencing. Immunoblot analysis using total cell lysates and the anti-HTT D7F7 antibody (Cell Signaling, 5656, 1:1000) confirmed the increased molecular weight of HTT. Additionally, confocal microscopy using anti-mNeonGreen (Chromotek, 32F6, 1:500) confirmed the expression of the HTT-mNG fusion protein.

Mouse ST*Hdh*^Q7^-*Hap40*KO-clone21 (ST*Hdh*^Q7^-*Hap40*KO) and ST*Hdh*^Q111^-*Hap40*KO-clone18 (ST*Hdh*^Q111^-*Hap40*KO) cell lines were generated using CRISPR-Cas9 genome editing. The strategy for genetic editing employed the use of two sgRNA sequences expressed from different pSpCas9(BB)-2A-Puro plasmids. The two gRNA sequences were: TCTGCGTCCTCCTTGGGCGG and CTTCTTGGCACGCTATCGGC. Each sgRNA induces a double-strand break at a distinct position near the start of the mouse *Hap40* gene, producing a deletion of up to 50 base pairs that results in a frameshift and a premature stop codon. The transfection and selection workflow were similar as mentioned above. Here, 150,000 cells were seeded into a six-well plate and 1 µg of each plasmid was transfected into cells using FuGENE transfection reagent at a 3:1 reagent µL:µg DNA ratio. Selection was performed with puromycin (2 µg/mL for STHdh^Q7^ and 4 µg/mL for ST*Hdh*^Q111^). Single-cell colonies were screened by PCR for the 50 bp deletion using the knockout primer pair CATTTGCGTCACTCAAGCCC (forward) and CAAGAAGTCCCCAGCCTCTG (reverse). An unedited clone yielded an amplified fragment of approximately 361 bp, whereas a successfully edited knockout clone produced no detectable product. Clones confirmed as knockouts were subjected to a second PCR with primers flanking the edited region (forward CGGAAGCGGAGGTGATGAAT, reverse CACGTTCGGCTTCCTCAAGA). Sanger sequencing of this amplicon verified the intended deletion and demonstrated a frameshift mutation that generates a premature stop codon. Chromatogram analysis was performed using Benchling.com software. Finally, immunoblot analysis of total cell lysates with the anti-HAP40 antibody (Atlas, HPA046960,1:500) confirmed the loss of full-length HAP40 expression in the knockout clones.

### Immunofluorescence

HEK293 and mouse striatal cells were detached, counted, and then seeded on either uncoated coverslips (mouse striatal cells) or on fibronectin/poly-L-lysine-coated coverslips (HEK293). For coverslip coating, 12 mm coverslips in 24-well plates were washed once with PBS (Thermo Fisher Scientific, 14190-094) and then incubated with fibronectin (Sigma, F1141) and poly-L-lysine (Sigma, P8920), both diluted 1:100 in ddH2O, for four hours at 37°C. Following this incubation, the coverslips were washed once with ddH2O and once with PBS before the cells were seeded on top. HEK293 and mouse striatal cells were seeded between 20,000 to 25,000 cells per well. Cells were fixed 48 hours after seeding as follows: coverslips were washed once with PBS and incubated with 4% PFA (Thermo Fisher Scientific, 8908) for 10 minutes at room temperature. The PFA was removed, cells were washed 3x with PBS and then permeabilized using 0.01% Triton-X 100 (Sigma Aldrich, T9284) for 10 minutes at room temperature. After permeabilization, the cells were incubated with Hoechst (Thermo Fisher Scientific, H3570) (1:10,000 in PBS) for 2 to 5 minutes at room temperature. After washing the cells, the coverslips were incubated for one hour at room temperature with 3% bovine serum albumin (BSA) (Sigma, A3059-10G) in PBS for blocking. After blocking, the cells were incubated overnight at 4°C with primary antibodies against the proteins of interest diluted in 3% BSA: rabbit anti-HAP40 (Atlas, HPA046960,1:500), rabbit anti-LC3B (Abcam, ab192890, 1:500), mouse anti-mNeonGreen (Chromotek, 32F6, 1:500), anti-HTT MAB2166 antibody (Millipore, MAB2166, 1:500). The next day, primary antibodies were removed, cells were washed 5x with the 3% BSA solution and then incubated with the respective secondary antibodies Alex Fluor 488 (Thermo Fisher Scientific, A11008 or A11001) or Alexa Fluor 647 (Thermo Fisher Scientific, A21245 or A21236) diluted 1:500 in PBS for two hours at room temperature. The coverslips were washed 5x with PBS and then mounted on microscopy slides using DAKO mounting medium (Agilent technologies, S302380-2) and left to dry overnight at room temperature before imaging.

### Microscopy

HEK293 and mouse striatal cells were imaged using a Leica TCS SP8 WLL confocal microscope. All samples were imaged using the 63x oil immersion objective. Images were taken as Z-stacks with an average of 15-25 steps and a system-optimized Z-step size. To capture images from samples containing Hoechst and the secondary antibodies Alex Fluor 488 and/or Alexa Fluor 647, the following lasers were used: diode laser (405 nm), argon-ion laser (488 nm), and HeNe laser (633 nm), respectively. Imaging the mNG signal in HEK293-HTT-mNG cells was done using the argon-ion laser for fluorophore excitation and the emission was captured at 500-570 nm.

### Image Analysis Colocalization analysis

All images within a specific imaging experiment for HEK293 were processed with the same parameters using the Fiji software(85), unless otherwise stated. Channels were first split and then each channel median filters were applied, and background signals were reduced. The channels were then merged again, and a Z-project was created using the maximum intensities of all Z positions. Look up tables were assigned to each channel and images were analyzed. The co-localization of the signals from the green (HTT-mNG) and red (HAP40) channels was analyzed qualitatively and quantitively. For the former, a line scan analysis was done by drawing a line on the image going across the signals of interest and then obtaining the plot profile from each channel. The values from all three channels were then overlapped by plotting the gray values of the pixels against the distance in µm on the line. Quantitative analysis of colocalization was done by calculating the Pearson and Manders (M1 and M2) coefficients using the JACoP plugin in Fiji(86). After assigning the red and green channels to images A and B, the threshold was set for each channel separately according to the signal intensities. The coefficients obtained from this analysis were plotted on a bar graph using GraphPad Prism.

### LC3B puncta analysis

All images taken of striatal cells were processed in the same way using the Fiji software, unless otherwise stated. First, median filters were manually applied to the split channels, and the background signals were reduced. The channels were then merged again, and a Z-project was created using the maximum intensities of all Z positions. Look up tables were assigned to each channel and then the images were analyzed. For LC3B puncta count and size analysis, single cells were cropped out of images and analyzed separately to obtain the number and size of LC3B puncta per cell. The signal threshold for the LC3B channel was set manually according to the observed signals, then using the “Analyze particles” option in Fiji the number and area of the puncta were measured. For particle analysis, the size in micron^2^ was set to 0-infinity, the circularity was set to 0.0-1.0, and particles touching the borders of the image were excluded by selecting the option “exclude on edges”. The average count and size of puncta per cell were calculated and plotted on bar graphs using GraphPad Prism. Statistical analysis using a one-way ANOVA was also performed using GraphPad Prism.

### Cell treatments

Striatal cells were plated in 6-well dishes at a density of 200,000-300,000 cells per well in 2 mL of complete medium and incubated overnight at 33°C. The following day, drug treatment, 100 mM Bafilomycin A1 solubilized in DMSO (Cell Signaling, 54645S) or 5 µg/mL Brefeldin A (Cell Signaling, 9972), was applied in complete DMEM or Opti-MEM (secretion experiments) medium for the indicated times in figure legends.

### Viability assay

Striatal cells were seeded into a transparent 96-well plate at a density of 10,000 cells per well in 100 µL of complete medium. The next day, the number of cells was determined by staining cells with Hoechst 33342 (Thermo Fisher Scientific, H3570) at final concentration of 5 µg/mL in complete medium for 15 min at 33°C. Afterwards, wells were washed with PBS and imaged using a Tecan Spark Cyto device. Next, the CellTiter-Glo 2.0 cell viability assay (Promega, G924) was used by applying 25 µL of CellTiter-Glo per well. Plate was mixed using an orbital plate shaker: 2 min, 700 rpm and incubated for 10 min at room temperature. Total luminescence was measured with a Tecan Spark Cyto device with an integration time of 0.25-1 second per well.

### Protein secretion assay and trichloroacetic acid precipitation

Striatal cells were seeded into a 6-well plate at a density of 200,000-300,000 cells per well in 2 mL of complete medium and incubated overnight at 33°C. The following day, cells were washed 3x with warm PBS and 1.5 mL of warm Opti-MEM medium was added to each well. At this point, for certain experiments drug treatment was performed (see cell treatments), otherwise plates were incubated at 33°C for 24 hr. After this incubation period, all conditioned medium was collected in 2 mL tubes and placed on ice. Cell pellets were collected by trypsinization, neutralization with medium, and washed with PBS before freezing pellets. For these steps, centrifugation was performed at 700 x g for 5 min at 4°C. Collected conditioned medium was first centrifuged at 700 x g for 10 min at 4°C and supernatant transfer to a new 2 mL tube. Samples were centrifuged again at 2,000 x g for 20 min at 4°C and 1,370 µL was transferred to a new 2 mL tube. Proteins in precleared conditioned medium were precipitated with addition of 151 µL of 100% TCA. Samples were inverted and incubated for 10 min on ice. Afterwards, 500 µL of ice-cold 10% TCA was added, inverted to mix, and incubated for 20 min on ice. Samples were then centrifuged at 20,000 x g for 10 min at 4°C. Supernatant was carefully removed and samples incubated with 1 ml ice-cold acetone overnight at −20°C. The next day, samples were centrifuged at 20,000 x g for 10 min at 4°C and acetone carefully removed. Samples were left to air-dry before addition of 12 uL of sample loading buffer (100 mM DTT, 1x NuPAGE LDS sample buffer, diluted with TNT-lysis buffer). In parallel, the respective cell pellets were lyzed in 15 µL of TNT-lysis buffer (20 mM Tris-HCl pH 8.0, 150 mM NaCl, 1% Triton-X). Total cell lysates and conditioned medium was subjected to SDS-PAGE and immunoblotting together.

### Proteasomal activity assay

Striatal cells were seeded into a transparent 96-well plate at a density of 10,000 cells per well in 100 µL of complete medium and incubated overnight at 33°C. The next day, the number of cells per well was determined by staining a designated set of cells with Hoechst 33342 (Thermo Fisher Scientific, H3570) at final concentration of 5 µg/mL in complete medium for 15 min at 33°C. Afterwards, wells were washed with PBS and imaged using a Tecan Spark Cyto device, eight technical replicates were assessed per genotype to obtain a mean value. Next, the 20S Proteasome Assay Kit (Cayman Chemical, Cay10008041-1) was used for assessing proteasome activity. First, the plate was centrifuged at 500 x g for 5 min, medium was discarded from a separate designated set of wells, and 200 µL of 20S proteasome assay buffer was added to each well. Then, plate was centrifuged at 500 x g for 5 min, buffer was discarded, and 100 µL of 20S proteasome assay lysis buffer was added. Plate was incubated in an orbital shaker at 300 rpm for 30 min at room temperature. Afterwards, plate was centrifuged at 1000 x g for 10 min at room temperature. Then 90 µL of supernatant was transferred to a clear bottom black 96-well plate. Next, 40x substrate was diluted to 1x using assay buffer and 10 µL of solution was added to each well. Plate was incubated in an orbital plate shaker set to 300 rpm for one hour at 37°C. Fluorescence readout was performed with a Tecan Spark Cyto using the following settings: fluorescent intensity; Ex/360 nm, Em/480 nm.

### Lysosomal activity assay

Striatal cells were seeded in 6-well plates at a density of 200,000-300,000 cells per well in 2 mL of complete medium and incubated overnight at 33°C. The next day the medium was replaced with 1 mL of pre-warmed DMEM supplemented with 0.5% FBS, 1x Pen/Strep, and 15 µL of the self-quenched substrate from the Abcam Lysosomal Intracellular Activity Assay Kit (ab234622); all solutions were protected from light. Labeled cells were incubated for 1 hour at 33°C, then detached with trypsin, examined microscopically to confirm a single-cell suspension, and neutralized with complete medium. The cell suspension was transferred to a 1.5 mL microcentrifuge tube, gently resuspended with a 1 mL pipette tip to break up any remaining clumps and centrifuged at 700 x g for 3 min. The pellet was washed twice with 1 mL FACS buffer (PBS with 1% FBS), repeating the centrifugation step each time. Finally, the cells were resuspended in 400 µL cold FACS buffer and kept on ice until flow-cytometric acquisition.

### Autophagic vacuoles labelling

Striatal cells in logarithmic growth were allowed to reach 70-80% confluence. They were then washed with PBS, detached with trypsin, and resuspended in complete medium. After cell counting, 200,000 viable cells were allocated to each condition (treated and untreated). The cells were pelleted at 700 x g for 3 min and washed with phenol-free DMEM containing 5% FBS. A labeling mixture was prepared by adding 1 µL of the green reagent from the Autophagy Detection Kit (Abcam 139484) to 1 mL of phenol-free DMEM with 5% FBS. The cell pellets were resuspended in 250 µL of labeling solution or just medium (FACS negative control) and incubated for 30 min at 33°C, protected from light. Following incubation, the samples were centrifuged again at 700 x g for 3 min and washed twice with ice-cold FACS buffer. The final pellet was resuspended in 400 µL of cold FACS buffer and kept on ice until flow-cytometric acquisition.

### Flow cytometric analysis

Samples were analyzed using a BD LSR Fortessa cell analyzer equipped with 488 nm (530 / 30 BP) and 561 nm (710 /50 BP) lasers. Initial gating used forward-scatter versus side-scatter (FSC-A vs SSC-A) to isolate the cell population, and doublets were excluded by plotting FSC-H vs FSC-W followed by SSC-H vs SSC-W. The final gated population was evaluated on the 488 nm versus 561 nm channels. Untreated cells served as negative controls to define background fluorescence and set the final gates. Data acquisition was performed with FACSDiva software, collecting at least 10,000 live events per sample. Post-acquisition analysis was carried out in FlowJo v10 using the same gating strategy to obtain mean fluorescence intensity values, and statistical comparisons were made with GraphPad Prism v9.

### Western blots

Cell pellets were lysed in 30 to 50 µL HEPES lysis buffer (50 mM HEPES pH 7.0, 150 mM sodium chloride, 10% glycerol, 1% NP-40, 20 mM NaF, 1.5 mM MgCl_2_, 1 mM EDTA, 1 mM PMSF, 0.5% sodium deoxycholate, 1x Benzonase, 1x cOmplete EDTA-free protease inhibitor cocktail (Merck, 5056189001)) for 30 min on ice. Lysates were centrifuged at 14,000 rpm for 10 min at 4°C and supernatants collected. For experiments where detection for LC3B was required, samples were lysed for 5 min on ice in LC3-lysis buffer containing Dulbecco’s phosphate-buffered saline, 2% Triton-X, 1x cOmplete EDTA-free protease inhibitor cocktail and cleared by centrifugation at 16,000 x g for 10 min at 4°C. Samples treated with LC3-lysis buffer were lysed and subjected to SDS-PAGE on the same day. Protein concentrations were determined by BCA assay (Thermo Fisher Scientific, 23227) and 20-25 µg total protein was combined with 50 mM DTT and 1x NuPAGE LDS sample buffer, followed by 5 min at 95°C. Proteins and protein standards (Thermo Fisher Scientific, LC5925) were separated by SDS-PAGE using a NuPAGE 4-12% Bis-Tris gel with 1x MES buffer and wet transferred onto nitrocellulose membranes (Cytiva, 10600002) using transfer buffer (25 mM Tris, 192 mM glycine, 20% methanol) at 30V for two hours. For certain experiments, SDS-PAGE was performed using 3-8% Tris-Acetate gel with 1x Tris-Acetate SDS running buffer (Thermo Fisher Scientific, LA0041) and using the HiMark pre-stained protein standard (Thermo Fisher Scientific, LC5699). After transfer, membranes were stained with No-Stain protein labelling reagent (Thermo Fisher Scientific, A44717) following manufacturer’s instructions before blocking. Membranes were then blocked for one hour in 3% milk in PBS with 0.05% Tween. The following primary antibodies were applied overnight at 4°C: rabbit anti-HAP40 (Atlas Antibodies, HPA046960, 1:1000), anti-HTT D7F7 antibody (Cell Signaling, 5656, 1:1000), anti-HTT MAB2166 antibody (Millipore, MAB2166, 1:500), anti-HTT MW1 antibody (DSHB, MW1, 1:500), mouse anti-SQSTM1(p62) (abcam, ab56416, 1:1000), rabbit anti-LC3B (abcam, ab192890, 1:1000), rabbit anti-UBB (PTG, 10201-2-AP, 1:1000), rabbit anti-Cathepsin D (Cell Signaling 69854, 1:1000), rabbit anti-STX17 (PTG 17815-1-AP, 1:1000), mouse anti-FLAG (Sigma, F3165, 1:1000), mouse anti-mNeonGreen (Chromotek, 32F6, 1:200), rabbit anti-GFP (Abcam, ab290, 1:2500), mouse anti-c-Myc (Merck, M4439, 1:2000), mouse anti-Actin (abcam, ab8224, 1:1000), mouse anti-Tubulin (Sigma, T6074, 1:80,000), mouse anti-TIM23 (BD Biosciences, 611223, 1:1000), and mouse anti-GAPDH (Santa Cruz, sc-47724, 1:1000). The following POD-conjugated secondary antibodies diluted 1:2000 in 3% milk-PBS with 0.05% Tween and applied for one hour at room temperature: goat anti-Rabbit IgG peroxidase (Sigma, A0545), goat anti-mouse IgG peroxidase (Sigma, A0168). In certain cases, the following secondary antibody was applied at 1:2000 for one hour at room temperature in 1x fluorescence blocking solution (Thermo Fisher Scientific, 37565): IRDye 800CW Donkey anti-Rabbit IgG (Li-Cor, NC0964679) or IRDye 680RD Donkey anti-Mouse IgG (Li-Cor, NC0963034). Each membrane was incubated with WesternBright Quantum (advanstar, K-12042- D20) solution for two minutes, followed by acquisition of a chemiluminescence image using an iBright imaging system (Thermo Fisher Scientific). Band intensities were quantified using Fiji (ImageJ) and normalized to respective loading control within each corresponding lane. Statistical analysis was performed using Prism v9 software.

### Co-Immunoprecipitation

Cell pellets were lysed in IP lysis buffer (25 mM Tris pH 7.5, 150 mM NaCl, 5% glycerol, 1% IGPAL (NP-40),1x MS-SAFE protease, and 1x Phosphates inhibitor) for 30 min on ice. Lysates were centrifuged at 14,000 rpm for 10 min at 4°C and supernatants collected. Mouse brain tissue (400 mg) was homogenized using a Precellys device (5000 rpm, 2x 20 sec, break: 15 sec) with 1 mL ice-cold 50 mM Tris pH 7.5. The volume of homogenates was adjusted to a 10-fold excess with 1x RIPA Buffer (50 mM Tris pH 7.5, 150 mM NaCl, 1% Triton X-100, 0.1% SDS, 0.5% sodium deoxycholate, EDTA-free protease inhibitor and Benzonase 0.25 U/μL) and incubated for one hour on a rotating wheel at 4°C. Protein concentrations were determined by BCA assay (Thermo Fisher Scientific, 23227) and about one mg of lysate was used per co-IP experiment. Immunoprecipitation experiments were performed using the Pierce Crosslink Immunoprecipitation kit (Thermo Fisher Scientific, 26147) in combination with Dynabeads Protein G (Thermo Fisher Scientific, 10004D) following manufacturer’s instructions. Dynabeads were resuspended by vortexing for 30 seconds and for each co-IP reaction 12.5 µL of beads was transferred to an individual 2 mL LoBind microtube. Beads were placed on the magnetic stand and storage liquid was discarded, followed by two washing steps with 200 µL 1x coupling buffer. For antibody crosslinking to protein G, beads were resuspended in 100 µL 1x coupling buffer together with respective amount of antibody as determined in antibody-bead saturation experiments (anti-HTT D7F7 antibody, Cell Signaling, 5656, 7 µL) and incubated at room temperature under constant agitation for 60 minutes. Afterwards, beads were briefly pulse-centrifuged up to 500 x g, followed by magnetizing beads and removal of supernatant. Beads were then washed twice with 200 µL 1x coupling buffer, magnetized, and supernatant was discarded. Next, 2.5 µL of 20x coupling buffer, 9 µL of 2.5 mM DSS in DMSO, and 38.5 µL MiliQ water was added to each bead preparation and placed under constant agitation for 60 min at room temperature. Beads were then magnetized, supernatant was removed, and beads were washed with 3x 500 µL elution buffer. Beads were then washed 3x with 500 µL lysis buffer. Then, lysis buffer was removed, and 1 mg of cell lysate in a final volume of 500 µL lysis buffer was added to beads. A volume corresponding to 25 µg of protein was set aside from the cell lysate and used as input sample downstream for western blotting. Samples were incubated overnight at 4°C on a rotating wheel for gentle end-over-end mixing. Afterwards, beads were magnetized and washed 2x with 500 µL wash buffer (0.05 % IGPAL (NP-40), 50 mM Tris pH 7.5, 150 mM NaCl, 5% glycerol) followed by 2x washing with 500 µL IGPAL-free wash buffer. For antigen elution, beads were resuspended in 24 µL 1x LDS sample buffer containing 100 mM DTT and incubated at 95°C for 10 minutes. Beads were then magnetized and supernatant was collected. Eluate samples were used immediately for SDS-PAGE or stored at −20°C.

### Size Exclusion Chromatography (SEC)

HEK293 cells were prepared in the following matter. A pellet of 80 million cells was resuspended in 800 µL lysis buffer containing HEPES, 150 mM NaCl, 1 mM EDTA, 1 mM MgCl_2_, 1 mM DTT, 0.1% NP40, protease inhibitor cocktail (cOmplete, EDTA-free; Roche, 05056489001) and phosphatase inhibitor (Pierce, A32957). Cell suspensions were mixed by pipetting up and down, and by vortexing. Lysis was supported by freezing cell suspensions on dry ice for 5 min, followed by thawing on ice. Then 20 U benzonase (1:10,000, Merck, 1016540001) were added and lysates incubated for 30 min on ice. Lysates were centrifuged at 10,000 x g for 15 min at 4°C. Mouse brains (12-month-old C57BL/6 wild-type) were prepared in the following matter. Brain samples (5 mg) were lysed utilizing a manual homogenizer in lysis buffer (50 mM HEPES pH 7.4,150 mM NaCl, 1.5 mM MgCl_2_,1 mM EDTA, 1 mM DTT, protease inhibitor cocktail, Benzonase (1:10,000). Volume of lysis buffer used was 10x the weight of tissue. Lysates were centrifuged at 15,000 rpm for 15 min at 4°C. For both cells and mouse brain lysates, supernatant was transferred to a new tube and protein concentration was determined using BCA Protein Assay (Pierce, 23228).

For SEC, an Äkta Purifier and a Superose 6 Increase 10/300 GL column (Cytiva, GE29-0915-96) were used. The column was equilibrated using lysis buffer (without phosphatase inhibitors). For calibration of column, high and low molecular weight kits (Cytiva, 28403842 and 28403841) were used. Supernatant of cell lysate (2.5 mg protein) was loaded on the column and fractionated. Fractions covering a wide molecular weight range, starting from void volume (Blue Dextran) and extending down to 13 kDa, were collected. Two fractions were always pooled. For protein precipitation, a fourfold volume of cold-acetone (99.7%) was added to the pooled fractions, mixed and incubated overnight at −20°C. The next day, proteins were precipitated by centrifugation at 3,428 x g at 4°C for 45 min. The supernatant (∼95%) was carefully taken off and disposed. The residual solution was centrifuged again at 3,428 x g at 4°C for 5 min. The remaining supernatant was carefully removed without disturbing the protein pellet and disposed. The pellet was dried for 5 min, resuspended in 100 µL gel loading buffer (1x NuPage LDS sample buffer, 50 mM DTT) and heat-denatured at 95°C for 5 min. Samples were subjected to SDS-PAGE using NuPAGE Novex 4-12% Bis-Tris gels (Thermo Fisher Scientific, NP0329) followed by transfer of proteins onto a nitrocellulose membrane (0.45 µm; Cytiva, 10600002) using a wet blotting system (BioRad). Membranes were blocked and washed as described in Western blot section. Immunoblotting using anti-HTT D7F7 (Cell Signaling, 5656S, 1:1000) and anti-HAP40 (Atlas Antibody, HPA 046960, 1:500) antibodies in 3% milk/PBS was performed, followed by labelling with secondary peroxidase-conjugated anti-rabbit antibody (1:2000) for one hour. Membranes were incubated with WesternBrightTM Quantum (advanstar, K-12042-D20) solution and acquisition of a chemiluminescence image was performed using an iBright imaging system (Thermo Fisher Scientific).

### Plasmids

pDEST26-cmyc-HTTQ23 (full-length) and pDEST26-cmyc-HTTQ145 (full-length) were subcloned from pDONR221-HTTQ23 (human) or pDONR221-HTTQ145 (human) into pDEST26-cmyc-Gateway (GW), a gift from Matthias Selbach. pDEST26-FLAG-HA-Human HAP40 was subcloned from pDONR221-Human HAP40 into pDEST26-FLAG-HA-GW, a gift from Matthias Selbach. pDEST-CMV mCherry-GFP-LC3B WT was a gift from Robin Ketteler (Addgene plasmid #123230)(87). pSpCas9(BB)-2A-Puro-sgRNA Human HTT (3’), pSpCas9(BB)-2A-Puro-sgRNA Human F8A1 (5’), pSpCas9(BB)-2A-Puro-sgRNA#1 Mouse F8A1, and pSpCas9(BB)-2A-Puro-sgRNA#2 Mouse F8A1 were generated by ligating annealed sense and anti-sense oligos into BbsI-digested pSpCas9(BB)-2A-Puro (PX459) V2.0 vector (Addgene plasmid #62988). pMK-Human HTT-mNeonGreen homology-directed repair (HDR) template was designed using Benchling.com software, in which the mNeonGreen gene is flanked (left and right) by 500 base pairs corresponding to the integration site located before the stop codon of the HTT gene. The HDR template was synthesized by gene synthesis and cloned into the pMK backbone (Thermo Fisher Scientific). pDONR221-human HAP40 mutants (M2, M3, M5) were generated via site-directed mutagenesis of the pDONR221-HAP40 (human) plasmid. Plasmids used for BRET assays were generated by shuttling open reading frames in the pDONR221 donor vector into destination vectors (pcDNA3.1-PA-mCit-GW, pcDNA3.1-GW-PA-mCit, pcDNA3.1-NL-GW, pcDNA3.1-GW-NL) as previously described(33). BRET control plasmids (pcDNA3.1-PA-NL, pcDNA3.1-NL, pcDNA3.1-PA-mCit, pcDNA3.1-PA-mCit-NL) were previously generated and described(33). Generation of HTT intramolecular BRET sensor was previously described(88). Protein purification expression constructs for full-length HTT and HAP40 were previously described(18, 89). pFBOH-MHL-HAP40-M2, -M3, and -M5 vectors were generated via site-directed mutagenesis of the pFBOH-MHL-HAP40 vector (Addgene plasmid #124060). The following primers were used for mutagenesis: for the M2 construct; FWD TCCCAAGAAGCTCTTTCTGCTGCTCCAG and REV AAAGAGC-TTCTTGGGAAGCTGGCCGCTGCTC; for the M3 construct, FWD TTCGTCTTCTTT-TTGTGGGTAGCCATGACCAAAG and REV ACAAAAAGAAGACGAAAGCCATCAAGTCGCTGCAG. All four mutagenic primers (M2_FWD, M2_REV, M3_FWD, M3_REV), were used to generate the M5 construct.

### Transfections

For overexpression studies, one million HEK293 cells or 300,000 mouse striatal cells in a final volume of 2 mL complete DMEM medium were reversed transfected in six-well plates with a transfection mix composed of two µg of DNA in a volume of 100 µL Opti-MEM medium (Thermo Fisher Scientific, 51985034) and 6 µL FuGENE transfection reagent (1:3 ratio) following manufacturer’s instructions. After 48 hours cells were dissociated using trypsin, washed with ice-cold PBS, pelleted and stored at −80°C until further processing for Western blotting.

### BRET screening

BRET assays were performed as described(33). Here, human cDNAs for full-length HTTQ23 and HAP40 were shuttled into destination vectors (pcDNA3.1-PA-mCit-GW, pcDNA3.1-GW-PA-mCit, pcDNA3.1-NL-GW, pcDNA3.1-GW-NL) using LR clonase technology according to manufacturer’s instructions (Thermo Fisher Scientific). HEK293 cells were reverse co-transfected with NanoLuc and mCitrine fusion constructs using linear polyethyleneimine (25 kDa, Polysciences, 23966) and incubated for 48 hours. Previously generated control vectors expressing only NanoLuc (pcDNA3.1-NL) or PA-mCitrine (pcDNA3.1-PA-mCit) were used as background controls. Additional readout controls included transfecting pcDNA3.1-PA-mCit-NL (tandem control), pcDNA3.1-PA-mCit + pcDNA3.1-NL (tag control), and pcDNA3.1-PA-NL (bleed-through control). Live cell BRET measurements were carried out in flat-bottom white 96-well plates (Greiner, 655983) with each tested interaction pair in triplicate. Infinite microplate readers M1000 or M1000Pro (Tecan) were used for the readout with the following settings: total luminescence, fluorescence of mCitrine recorded at Ex 500 nm/Em 530 nm, luminescence measured using blue (370–480 nm) and green (520–570 nm) band pass filters with 1,000 ms integration time. BRET ratios were calculated by dividing the background corrected intensity at 520–570 nm by the intensity obtained at 370–480 nm and subsequent donor bleed-through subtraction from NanoLuc only expressing wells (pcDNA-PA-NL), as previously described(33). Threshold for significance was a BRET ratio that was above the BRET ratio of both control interaction pairs.

### Intramolecular BRET assay

Screens were performed in 96-well microtiter plates at a density of 40,000 HEK293 cells per well. Cells were reverse co-transfected in duplicates using linear polyethylenimine with HTT intramolecular BRET sensor (150 ng, pcDNA-NL-HTT^Q23(2686-mCit)^) and increasing amounts of pcDNA-cmyc-HAP40^WT^ (0.08, 0.16, 0.31, 0.63, 1.25, 2.5, 5, 10 ng) or HAP40^M5^ plasmid (0.16, 0.31, 0.63, 1.25, 2.5, 5, 10, 20 ng). After 48 hours, mCitrine fluorescence (Ex/Em: 500 nm/530 nm; gain 80), total luminescence, and BRET (BLUE, 370–480 nm; GREEN 520–570 nm; both filters at 1,000 ms integration time) was assessed using the Infinite microplate reader M1000 (Tecan). BRET ratios were calculated by dividing the background corrected intensity at 520–570 nm by the intensity obtained at 370–480 nm and subsequent donor bleed-through subtraction from NanoLuc only expressing wells (pcDNA-PA-NL), as previously described(33). Statistical significance was determined via one-way ANOVA relative to HTT intramolecular BRET sensor alone.

### Protein expression and purification

Sf9 insect cells were used for HTT expression as previously described(18, 89). Briefly, P3 recombinant baculovirus was used to infect cells, after which point cells were grown at 37°C until 80-85% viability, typically within 72 hours post-infection. For HTTQ23-HAP40, a 1:1 HTTQ23-HAP40 baculovirus ratio was used for infection. Following growth of baculovirus infected Sf9 cells, the cells were harvested by centrifugation, lysed by two freeze-thaw cycles, and insoluble debris was pelleted by centrifugation at 29,416 x g for one hour. Soluble protein was purified by the supernatant by FLAG affinity chromatography, after which point the crude elution was pooled, concentrated, and polished by size-exclusion chromatography on a Superose 6 Increase 10/300 GL column (Cytiva Life Sciences) in gel filtration buffer (20 mM HEPES pH 7.4, 300 mM NaCl, 2.5% (v/v) glycerol, 1 mM TCEP). Purity of eluted fractions was assessed by SDS-PAGE, and fractions containing pure protein were pooled, concentrated, flash frozen in liquid N_2_, after which point aliquots were stored at −80°C until further use.

## Differential Scanning Fluorimetry

A Roche Applied Science Light Cycler 480 II was used for all differential scanning fluorimetry experiments. Protein samples were diluted in DSF buffer (20 mM HEPES pH 7.4, 150 mM NaCl, 1 mM TCEP, 5x Sypro Orange) to a final concentration of 100 µg/mL. 20 µL samples were plated in Roche LightCycler 384 well plates and protein unfolding was assessed by fluorescence measurement using 465 nm excitation and 580 nm emission filters over a 20-95°C temperature gradient (0.02°C/s ramp rate). Reactions were performed in triplicate. The temperature at which the first derivative of the melting curve reached a maximum was considered the melting temperature (Tm) for each protein complex studied.

### Proteomics Sample Preparation and Data Analysis

*HEK Immunoprecipitation-MS (IP-MS):* Sample preparation was conducted using on-bead tryptic digestion, adhering closely to the protocol established(90). Proteins immunoprecipitated from HEK293 cells were subjected to on-bead digestion and analyzed by label-free data-dependent acquisition (DDA) mass spectrometry. Beads were resuspended in 80 µL of digestion buffer (2 M urea, 50 mM Tris, pH 7.5) supplemented with 1 mM dithiothreitol (DTT) and 5 µg/mL sequencing-grade trypsin. The suspension was incubated for one hour at 25 °C with shaking at 1,000 rpm. The supernatant was collected, and the beads were washed twice with 60 µL of the same urea/Tris buffer. Supernatants and washes were combined (∼200 µL total), centrifuged at 5,000 x g, and transferred to a clean tube. Reduction was completed by adding DTT to a final concentration of 4 mM and incubating for 30 min at 25°C, 1,000 rpm. Alkylation was performed with 10 mM iodoacetamide (IAA) for 45 min at 25°C in the dark. An additional 0.5 µg of trypsin was added, and samples were digested overnight at 25°C with shaking at 700 rpm. Peptide clean-up was performed using in-house StageTips prepared with two C18 discs(91). Digests were acidified to pH < 3 with 1% formic acid. StageTips were conditioned with 100 µL of 90% acetonitrile / 0.1% formic acid (FA), then equilibrated with two washes of 100 µL 3% acetonitrile / 0.1% FA. Acidified samples were loaded, washed twice with 100 µL 3% acetonitrile / 0.1% FA, and eluted with 50 µL of 50% acetonitrile / 0.1% FA. All centrifugation steps were performed at 3,000 x g using benchtop spin-column adapters. Peptides were separated by nanoLC and analyzed on an Orbitrap Exploris 480 mass spectrometer (Thermo Fisher Scientific) coupled to a Vanquish Neo UHPLC system operated in nano-flow mode using a two hour gradient. Data were processed using MaxQuant v1.6.3.4 with iBAQ quantification and match-between-runs enabled. Searches were conducted against the Homo sapiens UniProt database (release 2018-07, including isoforms) supplemented with a custom HTT reference sequence (HTT.fasta, 21Q isoform). Protein groups identified as reverse hits, contaminants, or those identified solely by a modified peptide were excluded. Quantification was based on iBAQ values, with log2-transformation applied. Proteins were retained for downstream analysis if identified by at least two peptides and if at least one experimental group contained ≥4 valid values. Resulting data were median centered prior to statistical comparison.

*HEK Total Proteome (TMT):* HEK293 cell lysates were processed for Tandem Mass Tag (TMTpro) 16-plex labelling using an in-solution digestion workflow adapted from Mertins et al(92). Proteins were reduced with 5 mM dithiothreitol (DTT) for one hour at 37°C, followed by alkylation with 10 mM iodoacetamide (IAA) for 45 min at room temperature in the dark. Samples were diluted 1:4 with 50 mM Tris-HCl (pH 8.0) and digested sequentially with Lys-C (1:50, w/w) for two hours at room temperature, then with trypsin (1:50, w/w) overnight at room temperature. Digests were acidified to 1% formic acid (FA; pH ∼3), diluted to 1.5 mL with 0.1% FA, and cleared by centrifugation at 15,000 x g for 15 min. The supernatant was desalted using SepPak tC18 500 mg cartridges (Waters). For desalting, cartridges were conditioned with 3 mL acetonitrile (ACN) followed by 3 mL of 50% ACN / 0.1% FA, then equilibrated with four volumes of 3 mL 0.1% trifluoroacetic acid (TFA). Peptides were loaded, washed three times with 3 mL 0.1% TFA and once with 3 mL 1% FA, then eluted with 2x 1.5 mL of 50% ACN / 0.1% FA. The eluate was divided into 10 aliquots (∼100 µg each), dried by vacuum centrifugation, and stored at –20 °C. For TMT labelling, 100 µg of desalted peptide was resuspended in 20 µL of 100 mM HEPES (pH 8.0). TMTpro reagents (Thermo Fisher Scientific) were equilibrated to room temperature, reconstituted in anhydrous ACN (0.5 mg per 20 µL), and vortexed for 5 min. For each labelling reaction, 4 µL of TMTpro reagent was added to the peptide solution and incubated for one hour at room temperature. Reactions were quenched with 1 µL of 5% hydroxylamine for 15 min. Labelled peptides were pooled in equimolar amounts, dried by SpeedVac, reconstituted in 1 mL of 3% ACN / 0.1% FA, and desalted using SepPak tC18 200 mg cartridges (Waters). High-pH reversed-phase fractionation was carried out using an UltiMate 3000 HPLC system (Thermo Fisher Scientific), and samples were separated into 24 fractions using a gradient optimized for TMT-labelled peptides. Each fraction was dried and stored prior to LC-MS analysis. Peptides were analyzed on an Orbitrap Exploris 480 mass spectrometer (Thermo Fisher Scientific) coupled to an EASY-nLC 1200 system (Thermo Fisher Scientific) with a two hours LC gradient. Raw MS data were processed using MaxQuant v2.0.3.0(93). Spectra were searched against the Homo sapiens UniProt reference proteome (release 2018-07, including isoforms) combined with a database of common proteomics contaminants. Reporter ion intensities were log2-transformed and normalized using median–MAD scaling. Data were filtered to remove reverse hits, common contaminants, and identifications based solely on site-modified peptides. Protein groups were retained if supported by at least two peptides and if complete reporter ion intensities were available for all samples in the TMT plex. Differential abundance analysis for all three MS datasets was performed using limma-assisted moderated t-tests(94) with Benjamini–Hochberg false discovery rate (FDR) correction for multiple testing.

*ST Total Proteome (DIA):* An amount of 5 million striatal cells samples was lysed in 1x SDC buffer consisting of 1% (w/v) sodium deoxycholate (SDC; Sigma-Aldrich), 10 mM dithiothreitol (DTT; Sigma-Aldrich), 40 mM chloroacetamide (CAA; Sigma-Aldrich), and 100 mM Tris-HCl (pH 8.0). Lysis was carried out at 95°C for 10 min to ensure complete protein denaturation and simultaneous reduction and alkylation. Proteins were enzymatically digested overnight at 37°C using endopeptidase Lys-C (Wako) and sequencing-grade trypsin (Promega) at a protein-to-enzyme ratio of 50:1. Peptide clean-up was performed using the AssayMAP Bravo automated liquid handling platform (Agilent Technologies). Peptides were analyzed using a label-free data-independent acquisition (DIA) workflow on an Orbitrap Exploris 480 mass spectrometer (Thermo Fisher Scientific) coupled to a Vanquish Neo UHPLC system (Thermo Fisher Scientific) operating in nano-flow mode with a two hour reversed-phase LC gradient. Raw data were processed using DIA-NN v1.9(95), with searches conducted against the Mus musculus UniProt reference proteome (release 2022-03, including isoforms) supplemented with a custom database containing HTT and HAP40 sequences (mouse_HTT_HAP40.fasta). Protein-level identification thresholds were set at Protein.Q.value ≤ 0.01, Lib.Q.Value ≤ 0.01, and Lib.PG.Q.Value ≤ 0.01. Quantification was performed by aggregating precursor intensities using maximum selection across each Run, Protein Group, and Gene. Protein intensities were log2-transformed and quantified using the MaxLFQ algorithm(35). Proteins were retained for further analysis if supported by at least two peptides and if log2-transformed intensities were present in at least 75% of all samples. Missing values were imputed column-wise using a left-censored normal distribution with a randomized Gaussian downshift.

*Striatal cells Immunoprecipitation-MS (IP-MS)*. Following immunoprecipitation (1 mg input) on-bead digestion of the samples was performed in 6M Urea (100 mM ammonium bicarbonate, pH 8.0) for 2 hours at 37°C with 500 ng of Lys-C in the presence of 5 mM tris(2-carboxyethyl)phosphine (TCEP; Sigma-Aldrich) and 20 mM CAA. Samples were subsequently diluted to 1.6M Urea and further digested with 1000 ng of trypsin overnight at 37°C. The supernatant was quenched with TFA (1%) and the peptides cleaned using C18 columns (20 mg HNS S18V; The Nest Group) using standard techniques. Peptides were measured using a DIA workflow on an Orbitrap Exploris 480 mass spectrometer coupled to a Vanquish Neo UHPLC system operating in nano-flow mode utilizing a 60 min non-linear gradient. Data analysis was performed using Spectronaut 20 (Biognosys, 20.2.250922.92449)(96) against the canonical and reviewed Mus musculus database (release 2026-03; 17,246 entries) with default settings and cross-run normalization and imputation enabled. The data was also exported without imputation to filter for proteins that were only found in all biological replicates in at least one experimental group. Differential abundance analysis was performed as described above. For all MS datasets, differential abundance analysis was performed using limma-assisted moderated t-tests(94) with Benjamini–Hochberg false discovery rate (FDR) correction for multiple testing.

### RNA extraction

For quantitative real-time PCR and RNA-sequencing, RNA was purified from mouse striatal cells (70% sub-confluent T175 flask) using a RNeasy kit (Qiagen). DNA contamination was removed from purified RNA by using the RNase-Free DNase set (Qiagen). RNA concentration was determined by measuring the absorbance at 260 nm, and purity was assessed via the 260/280 nm ratio.

### Reverse Transcription and Quantitative Real-Time PCR

Quantitative real-time PCR was adapted as previously described(97). Single-stranded cDNA from total RNA was synthesized using the High-Capacity cDNA Reverse Transcription kit (Thermo Fisher Scientific, 4374967). For qRT-PCR, 50 ng of cDNA per well was amplified in a 10 µL reaction containing TaqMan Gene Expression Master Mix (Thermo Fisher Scientific) and a TaqMan probe specific for mouse *Htt* (Assay ID Mm01213820_m1; primers span exons 64-65). For normalization of data, an endogenous mouse *Actb* (Mm02619580_g1, Thermo Fisher Scientific) FAM/MGB labelled probe was used. Real-time PCR was performed using the ViiA 7 real-time PCR system (Applied Biosystems). Samples were measured in triplicates. Quantification was performed using the ΔCt met.

### RNA Sequencing

Total RNA samples were quantified using a Qubit Fluorometer, and RNA integrity was checked on a TapeStation (Agilent Technologies). Double-indexed stranded mRNA-Seq libraries were prepared using the Illumina Stranded mRNA Prep kit (50040534), starting from 500 ng of input material according to the manufacturer’s instructions. Libraries were equimolarly pooled based on Qubit concentration measurements and TapeStation size distributions. The loading concentration of the pool was determined using a qPCR assay (Roche, 7960573001). Libraries were then sequenced on the Illumina NovaSeq X Plus platform using PE100 sequencing mode, with a target of 50 million reads per library.

### RNA-Seq Data Processing and Differential Expression Analysis

RNA-Seq reads were aligned to the mouse genome (GRCm38, Ensembl release 87, version M12) using STAR(98) (v2.7.3a) with the following parameters: outSAMunmapped Within, outFilterType BySJout, outFilterMultimapNmax 20, alignSJoverhangMin 8, alignSJDBoverhangMin 1, outFilterMismatchNmax 999, outFilterMismatchNoverLmax 0.04, alignIntronMin 20, alignIntronMax 1000000, alignMatesGapMax 1000000. Read assignment to genes was performed with FeatureCounts(99) (v2.0.0) using -T 2 -t exon -g gene_id -s 2 -p. Differential expression analysis was conducted with DESeq2(100) (v1.38.0), excluding genes with fewer than five counts in at least three samples (**Table S3**).

### Principal Component Analysis (PCA)

Principal component analysis (PCA) was conducted in R (v4.5.0) using the prcomp function from the stats package to reduce the dimensionality of the dataset and identify the major axes of variation. Prior to PCA, expression values were log-transformed and scaled to have zero mean and unit variance across features. PCA plots were generated by visualizing the first two principal components, with samples colored by experimental group. For RNA-Seq expression profiles, PCA was applied to mouse striatal cell lines expressing either wild-type HTT (ST*Hdh*^Q7^) or mutant HTT (ST*Hdh*^Q111^), each with and without *Hap40* knockout.

### Gene Ontology (GO) and KEGG enrichment analyses

Gene Ontology enrichment analyses were performed in R (v4.5.0) using the enrichGO function from the clusterProfiler package(101) (v4.16.0) to identify overrepresented terms. Statistical significance was assessed using a hypergeometric test, and p-values were adjusted for multiple testing with the Benjamini–Hochberg false discovery rate (FDR) method. Terms with FDR ≤ 0.05 were considered significant. Proteins significantly enriched with HTTQ23-HAP40 complexes in WT cells or with apo-HTTQ23 in HAP40 KO cells, excluding those also enriched in HTT KO extracts, were analyzed. KEGG pathway enrichment analysis was carried out with shinyGO v0.8(102) using the KEGG database as the reference. Gene identifiers supplied as gene symbols were automatically mapped to Ensembl/STRING IDs via shinyGO’s built-in conversion utility. Parameters for analysis included: human background, FDR cutoff of 0.001, redundancy removed, background all protein-coding genes.

### Ingenuity Pathway Analysis (IPA)

Proteomic and transcriptomic datasets were analyzed using Ingenuity Pathway Analysis(103) to identify enriched canonical pathways and biological functions. The significance of each pathway was calculated in IPA using a right-tailed Fisher’s exact test, which estimates the probability that the observed association between the dataset and a given pathway occurs by chance. p-values were subsequently adjusted for multiple testing using the Benjamini–Hochberg false discovery rate (FDR) method. Only the most significant pathways after p-value correction were selected for reporting.

### CLEAR and lysosomal omics

For pathway-focused analyses, we used a curated list of genes associated with the Coordinated Lysosomal Expression and Regulation (CLEAR) network, obtained from Ingenuity Pathway Analysis (IPA) Database. In parallel, we used a reference list of lysosome-associated genes and proteins compiled from the literature(37, 38). In Figure 4D, transcriptomic data from ST*Hdh*^Q7^-*Hap40*KO and ST*Hdh*^Q111^-*Hap40*KO cells were intersected with the CLEAR gene list to identify significantly upregulated members of this regulatory network. In Figure 4E, differentially expressed genes from the same RNA-Seq analysis were filtered using the lysosomal reference list to highlight transcriptional changes in lysosome-associated genes. In Figure 4F, proteomic datasets from the same cell lines were compared against the lysosomal reference protein list to assess corresponding changes at the protein level.

## Acknowledgments

The authors thank Roxanne Maria Papawassiliou, Megan Bonsor, Anne Ast, and Adrian Marti Pastor for technical support. The authors thank the MDC/BIH Genomics Technology Platform at the Max Delbrück Center for Molecular Medicine in the Helmholtz Association and the Berlin Institute of Health (BIH), Berlin, Germany for technical support and assistance in this work. We thank the Advanced Light Microscopy Technology platform at the MDC Berlin for general and technical support. We are grateful to Dr. Hans-Peter Rahn and Kristin Rautenberg at the Flow Cytometry Technology Platform at MDC Berlin for operational support. E.E.W discloses funding support from CHDI foundation. E.S.R thanks the Huntington’s Disease Society of America for funding support through the Berman-Topper Family HD Career Development Fellowship.

## Author Contributions

Conceptualization, E.E.W., E.S.R.; Methodology, E.S.R., A.B., P.T., C.S., F.S., O.A., O.P., A.I., B.K., R.J.H., L.R., T.K., N.N., N.S., M.Z., S.G., S.B.; Formal Analysis, E.S.R., A.B., P.T., F.S., C.H., O.A., C.S., O.P., A.I., R.J.H.; Writing–Original Draft, E.E.W., E.S.R.; Writing Review & Editing, E.E.W., E.S.R., S.S.; Supervision, E.E.W., E.S.R., A.B., P.T., C.S., P.M., I.P.; Visualization, E.S.R., A.B., P.T., C.H., O.A., C.S., O.P., R.J.H.; Funding Acquisition, E.S.R., E.E.W.

---

**Figure S1:**
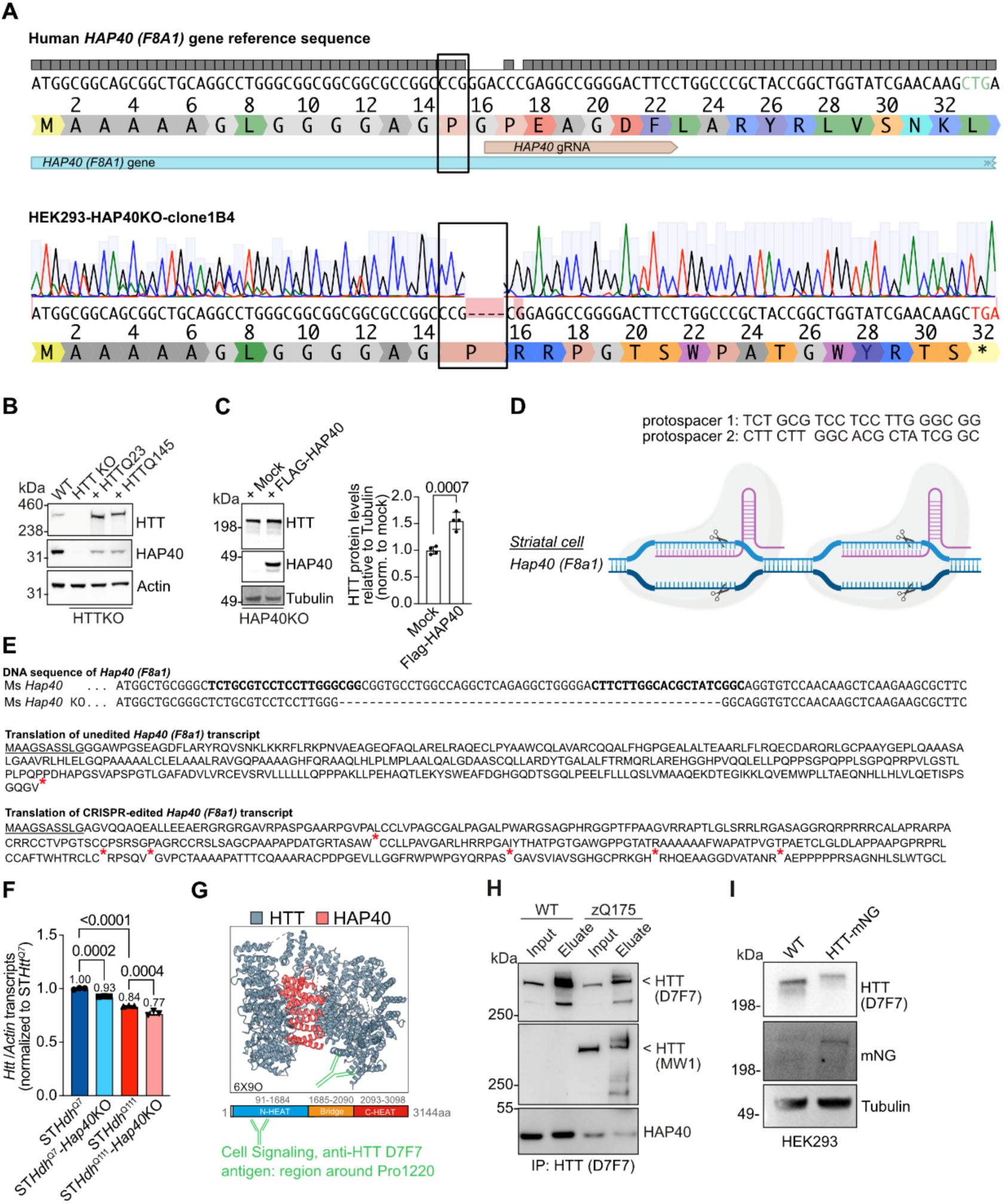
Characterization of cell lines and mouse brains. **A)** Sequence chromatogram of HEK293-HAP40KO-clone1B4 showing the CRISPR/Cas9-induced insertion (indel) mutation. The indel causes a frameshift, resulting in a premature stop codon in the *HAP40* (*F8A1*) gene. **B)** Representative immunoblot of HEK293 total cell lysates. HEK293-HTTKO cells were transfected for 48 hours with plasmids for expression of full-length cmyc-HTTQ23 or cmyc-HTTQ145. Samples were immunoblotted using anti-HTT MAB2166, anti-HAP40, and anti-beta Actin antibodies. **C)** Left, representative immunoblot of HEK293-HAP40KO total cell lysates. HEK293-HAP40KO cells were transfected for 48 hours with plasmids for expression of FLAG-HAP40 or mock (FLAG backbone plasmid). Samples were immunoblotted using anti-HTT (D7F7), anti-HAP40, and anti-tubulin antibodies. Right, quantification of the HTT protein levels relative to Tubulin. Data are presented as mean ± SD of four biological replicates. Statistical significance was assessed by student t-test**. D)** Schematic representation of the CRISPR/Cas9 editing strategy utilizing two sgRNAs to induce a 50 base pair deletion in the *Hap40* (*F8a1)* gene of mouse striatal cells. Sequence protospacer for each sgRNA is displayed. Created in BioRender. Wanker, E. (2026) https://BioRender.com/cr6sk7g. **E)** Top: DNA sequence alignment of wild-type and CRISPR-edited cells for the mouse *Hap40* (*F8a1)* gene. Dashed lines in the alignment indicate the absence of sequencing reads for the specific base. Translation of the *Hap40* (*F8a1)* transcript is shown for the wild-type (middle) and CRISPR-edited cells (bottom). Underlined amino acids represent the unaltered amino acid sequence relative to wild-type sequence and asterisk denotes the position of a stop codon. **F)** Analysis of steady-state *Htt* transcript levels in mouse striatal cells by qRT-PCR. Specific mouse probes for *Htt* mRNA and *Actin* mRNA were used, *Actin* mRNA was used as a reference gene. Data is expressed as means ± SD, n = 4-5 biological replicates. Statistical significance was determined via one-way ANOVA followed by Tukey’s multiple comparisons test. **G)** Top: Cryo-EM structure of HTT-HAP40 (6X9O) illustrating the general location of the anti-HTT (D7F7) target antigen. Bottom: schematic representation of HTT protein sequence, highlighting in N-HEAT, Bridge, and C-HEAT regions, along with the general position of the anti-HTT (D7F7) target antigen in relation to Proline 1220. **H)** Representative immunoblot of immunoprecipitations of HTT with anti-HTT (D7F7) antibody using mouse brains lysates from 12-month-old wild-type and zQ145 mice. IP fractions represent 5% input sample and 100% eluate. Samples were subjected to SDS-PAGE and immunoblotted for HTT using anti-HTT (MAB2166), anti-HTT (D7F7), anti-HTT (MW1, specific for HTT polyQ), and anti-HAP40 antibodies. Three independent experiments were performed. **I)** Representative immunoblot of HEK293 wild-type and HEK293-HTT-mNG knock-in total cell lysates. Membranes were probed with antibodies against anti-HTT (D7F7), anti-mNeonGreen (mNG), and anti-Tubulin antibody.

**Figure S2:**
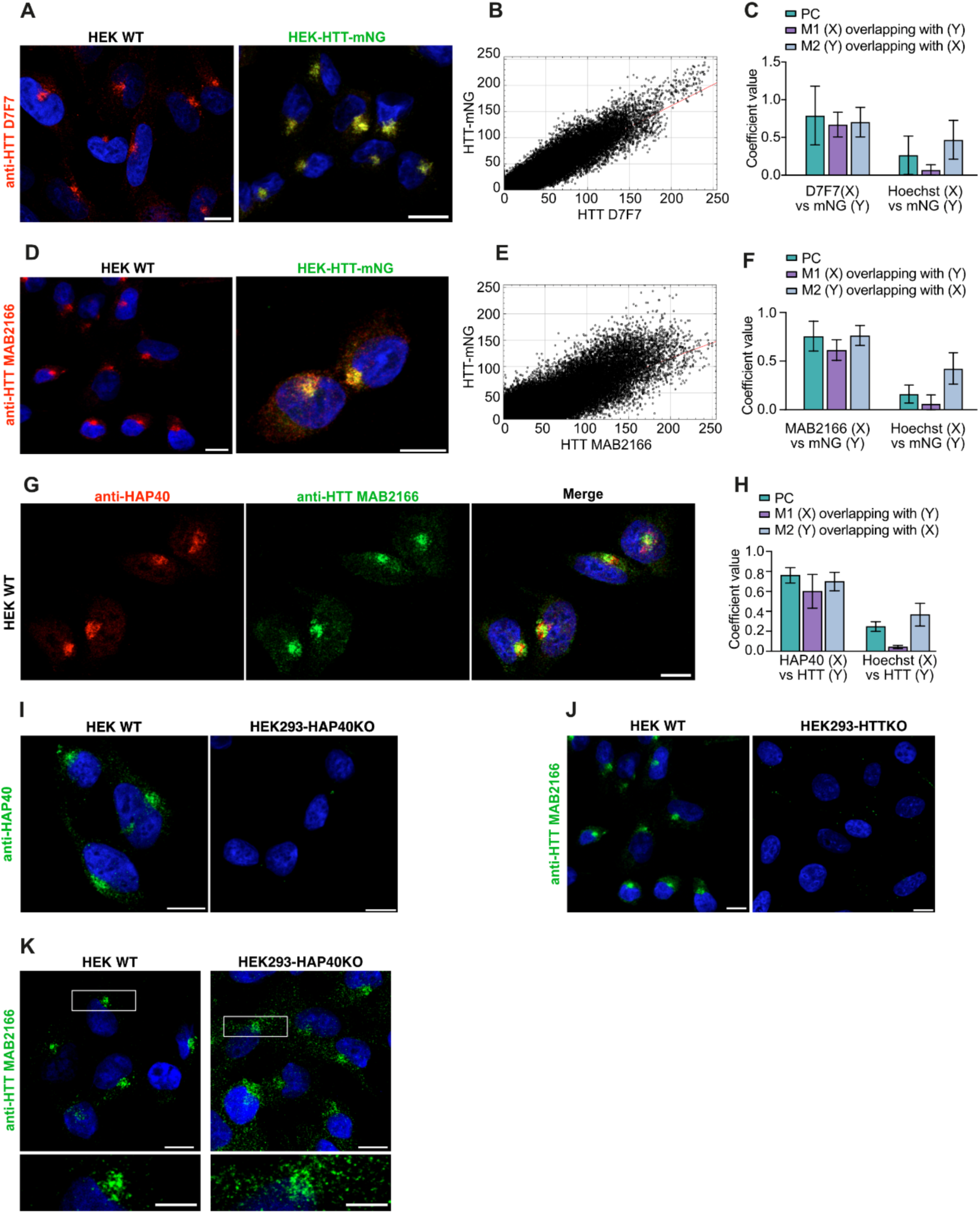
Control experiments related to HTT-HAP40 confocal imaging. **A)** Representative confocal images of HEK293 wild-type and HEK293-HTT-mNG cells immunostained for Huntingtin (HTT D7F7 antibody, red), mNeonGreen (green) and counterstained with Hoechst for nuclei (blue). **B)** Pixel intensity scatter plot showing the relationship between fluorescence intensities of Huntingtin (HTT D7F7 antibody, X-axis) and mNG (Y-axis). Each dot represents the intensity values of a single pixel in the two channels. These data were used to calculate Pearson’s correlation coefficients and Manders coefficients. **C)** Bar graph showing colocalization analysis between anti-HTT (D7F7) (X) and anti-mNG (Y) signals, as well as the colocalization of Hoechst (X) and HTT-mNG (Y) as a negative control. The analysis includes the average Pearson’s coefficient (PC) and Manders M1 (amount of signal from channel X overlapping with channel Y) and M2 (amount of signal from channel Y overlapping with channel X) coefficients. Data represent mean ± SD (n=3). **D)** Representative confocal images of HEK293 wild-type and HEK293-HTT-mNG cells immunostained for Huntingtin (HTT MAB2166 antibody, red), mNeonGreen (green) and counterstained with Hoechst for nuclei (blue). **E)** Pixel intensity scatter plot showing the relationship between fluorescence intensities of Huntingtin (HTT MAB2166 antibody, X-axis) and mNG (Y-axis). Each dot represents the intensity values of a single pixel in the two channels. These data were used to calculate Pearson’s correlation coefficients and Manders coefficients. **F)** Bar graph showing colocalization analysis assessing anti-HTT (MAB2166) (X) and anti-mNG (Y) signals, as well as the colocalization of Hoechst (X) and HTT-mNG (Y) as a negative control. The analysis includes the average Pearson’s coefficient (PC) and Manders M1 (amount of signal from channel X overlapping with channel Y) and M2 (amount of signal from channel Y overlapping with channel X) coefficients. Data represent mean ± SD (n=3). **G)** Representative confocal images of HEK293 wild-type cells immunostained for Huntingtin (HTT MAB2166 antibody, green), HAP40 (red) and counterstained with Hoechst for nuclei (blue). **H)** Bar graph showing colocalization analysis assessing anti-HAP40 (X) and anti-HTT (MAB2166) (Y) signals, as well as the colocalization of Hoechst (X) and anti-HTT (MAB2166) (Y) as a negative control. The analysis includes the average Pearson’s coefficient (PC) and Manders M1 (amount of signal from channel X overlapping with channel Y) and M2 (amount of signal from channel Y overlapping with channel X) coefficients. Data represent mean ± SD (n=3). **I)** Representative confocal images of HEK293-HAP40KO cells immunostained for HAP40 (green) and counterstained with Hoechst for nuclei (blue). **J)** Representative confocal images of HEK293-HTTKO cells immunostained for Huntingtin (HTT MAB2166 antibody, green), and counterstained with Hoechst for nuclei (blue). **K)** Representative confocal images of HEK293 wild-type and HEK293-HAP40KO cells immunostained for and anti-HTT (MAB2166, green) and counterstained with Hoechst for nuclei (blue). All images were acquired at 63x magnification. Scale bars represent 10 µm for all, expect (K) zoom in image represents 5 µm.

**Figure S3:**
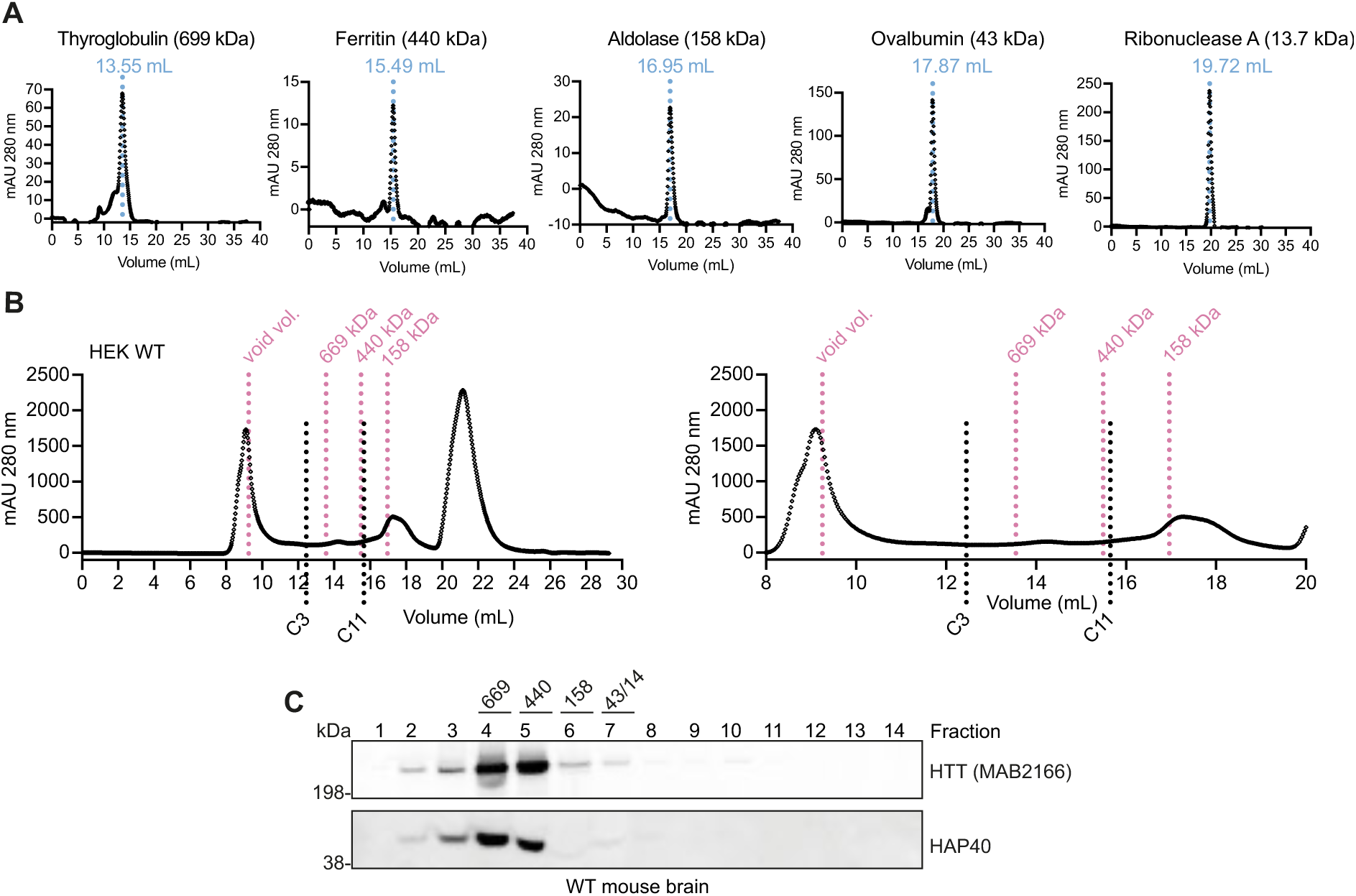
Size-exclusion chromatography analysis. **A)** Elution profiles of the molecular weight markers, including Thyroglobulin, Ferritin, Aldolase, Ovalbumin, and Ribonuclease A used for calibration of Superose 6 Increase 10/300 GL column. The x-axis represents the absorption measured at 280 nm, while the y-axis indicates the volume at which the marker protein elutes from the column. The elution volume is also noted above the peaks. **B)** Elution profile of HEK WT lysate analyzed by size-exclusion chromatography using a Superose 6 Increase column. The molecular weights of marker proteins and their corresponding elution volume (indicated by the magenta dotted line) are schematically shown at the top of the diagram. At the bottom, fractions C3 and C11 are indicated, eluting at 12.45 mL and 15.65 mL (black dotted line), respectively. A magnified section of the diagram is presented on the right side. **C)** Representative size-exclusion chromatography of total lysates from wild-type mouse brain. Fractions were pooled by combining three wells to obtain a total of 14 final fractions. Proteins were acetone-precipitated, and 50% of each fraction in 1x LDS was subjected to SDS-PAGE followed by immunoblotting using anti-HTT (MAB2166) and anti-HAP40 antibodies. Molecular weight markers are displayed above the fractions.

**Figure S4:**
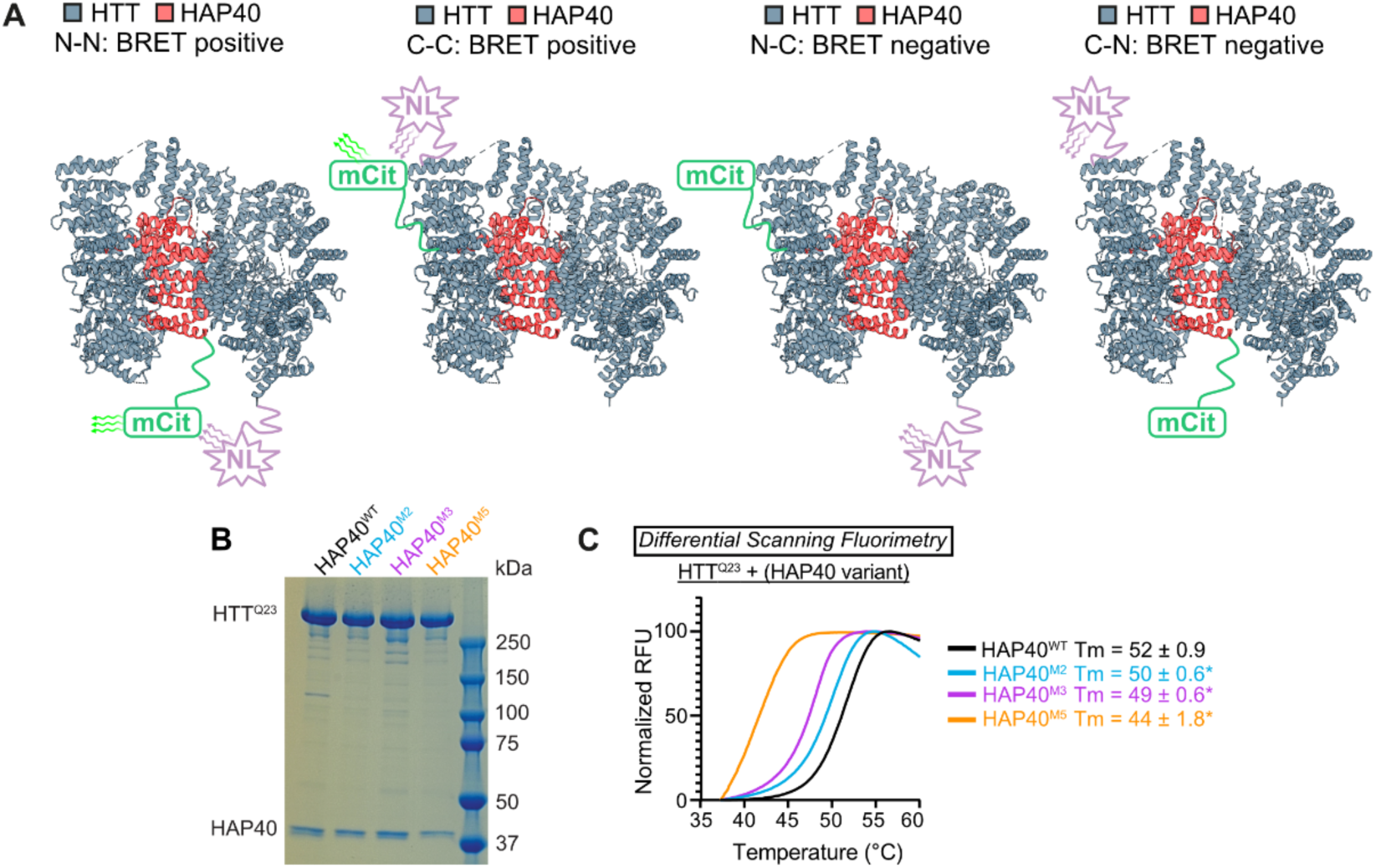
HTT intramolecular BRET senor and differential scanning fluorimetry. **A)** Cryo-EM structure of HTT-HAP40 (PDB: 6X9O) illustrating the different tagging orientations and BRET screening combinations. Screening with HTT and HAP40 tagged at the same termini (C–C or N–N) resulted in high BRET ratio values, whereas opposite orientations (N–C or C–N) showed non-significant BRET signals, likely due to the increased distance between donor and acceptor. **B)** Representative SDS-PAGE of purified full-length HTT^Q23^ in complex with wild-type HAP40 or the mutant variants HAP40^M2, M3, and^ ^M5^. **C)** Differential scanning fluorimetry profiles and calculated melting temperature (Tm) values for full-length HTT^Q23^ vs. wild-type HAP40 (black), HAP40^M2^ (teal), HAP40^M3^ (magenta), and HAP40^M5^ (orange) mutant variant. Melting temperatures were determined from the inflection point of curves obtained by fitting the data to the Boltzmann sigmoidal function. Significant (≥2 °C) destabilizations relative to the wild-type HAP40 are indicated with an asterisk. Data are presented as mean ± SD (n =3).

**Figure S5:**
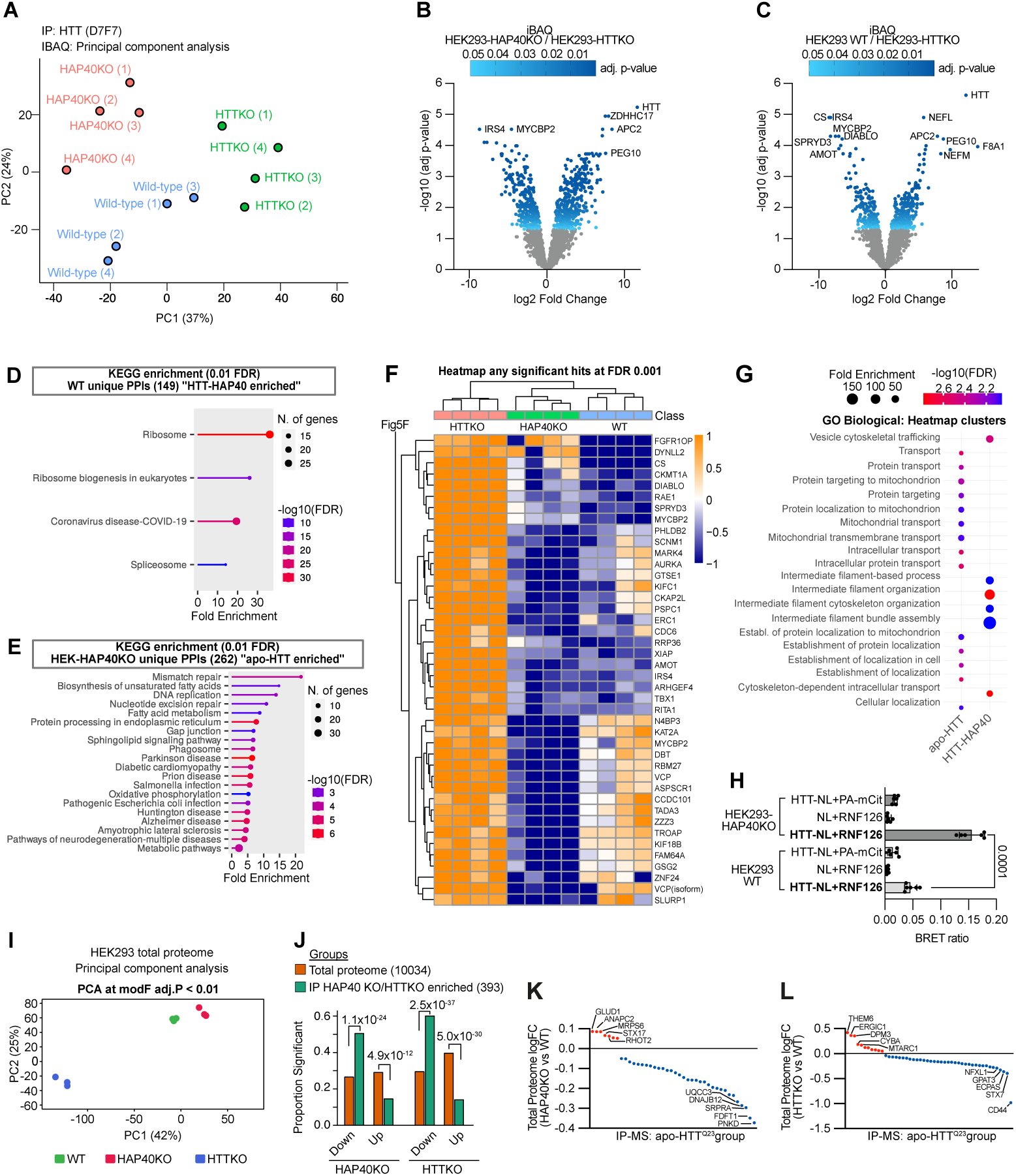
Multi-omics profiling of HTT-IP-MS and total proteome alterations in HEK293 wild-type, HTTKO, and HAP40KO cells. **A)** Principal component analysis of HEK293 HTT-IP-MS dataset. Points represent individual biological samples, colored by experimental group (wild-type, HTTKO, HAP40KO). The axes display PC1 and PC2 with the proportion of variance explained by each component indicated. **B)** Volcano plots of HTT-IP-MS iBAQ data comparing HEK293-HAP40KO vs HEK293-HTTKO and **C)** wild-type vs HEK293-HTTKO. The x-axes display log₂ fold changes, the y-axes shows -log₁₀ adjusted p-values (Benjamini–Hochberg). Shades of blue dots denote significantly enriched proteins (adjusted p ≤ 0.05); gray dots represent proteins that do not reach significance (adjusted p > 0.05). **D)** KEGG pathway enrichment for proteins significantly enriched in wild-type/HEK293-HTTKO and **E)** in HEK293-HAP40KO/HEK293-HTTKO HTT-IP-MS samples. Only pathways with a Benjamini–Hochberg FDR < 0.01 are shown. Plots show pathways ranked by fold enrichment and the number of constituent proteins is indicated by circle size and the -log₁₀ adjusted p-value in color scale. **F)** Heatmap related to Figure 5F of iBAQ intensities (row-wise Z-scaled) for proteins that passed a stringent enrichment threshold (FDR ≤ 0.001) across the three HEK293 cell lines. Hierarchical clustering was performed using Euclidean distance, revealing distinct enrichment patterns among the cell lines. **G)** Gene ontology (Biological Process) enrichment for the HTT-HAP40 enriched and apo-HTT enriched proteins. Each term is represented by a circle whose diameter represent the fold-enrichment value, while the color gradient reflects the Benjamini–Hochberg FDR. Only terms with FDR < 0.05 are displayed. **H)** BRET ratios of binary interactions between NanoLuc and mCitrine tagged full-length HTT^Q23^ and RNF126 and respective controls. Interactions were tested in HEK293 wild-type and HEK293-HAP40KO cells. Data are presented as mean ± SD, n=2, each with three technical replicates. **I)** Principal component analysis of HEK293 total proteome dataset. Points represent individual biological samples, colored by experimental group (wild-type, HTTKO, HAP40KO). The axes display PC1 and PC2 with the proportion of variance explained by each component indicated. **J)** Enrichment analysis of differentially regulated proteins. Two groups were examined: (i) the 393 IP-MS HAP40KO/HTTKO enriched proteins and (ii) the global HEK293 TMT-MS proteome in either HAP40KO or HTTKO cells. For each group, proteins were classified as significant or non-significant (adjusted p < 0.05) based on the TMT-MS data. Fisher’s exact test was applied to assess enrichment; p-values are displayed above the bars. **K)** LogFC values for the 62 proteins that clustered together in the hierarchical analysis shown in Figure 3F (apo-HTT enriched protein set) plotted against the total-proteome TMT-MS data for HEK293-HAP40KO versus wild-type. **L)** Same 62 protein set as in **K**, now plotted for the HEK293-HTTKO versus wild-type comparison.

**Figure S6:**
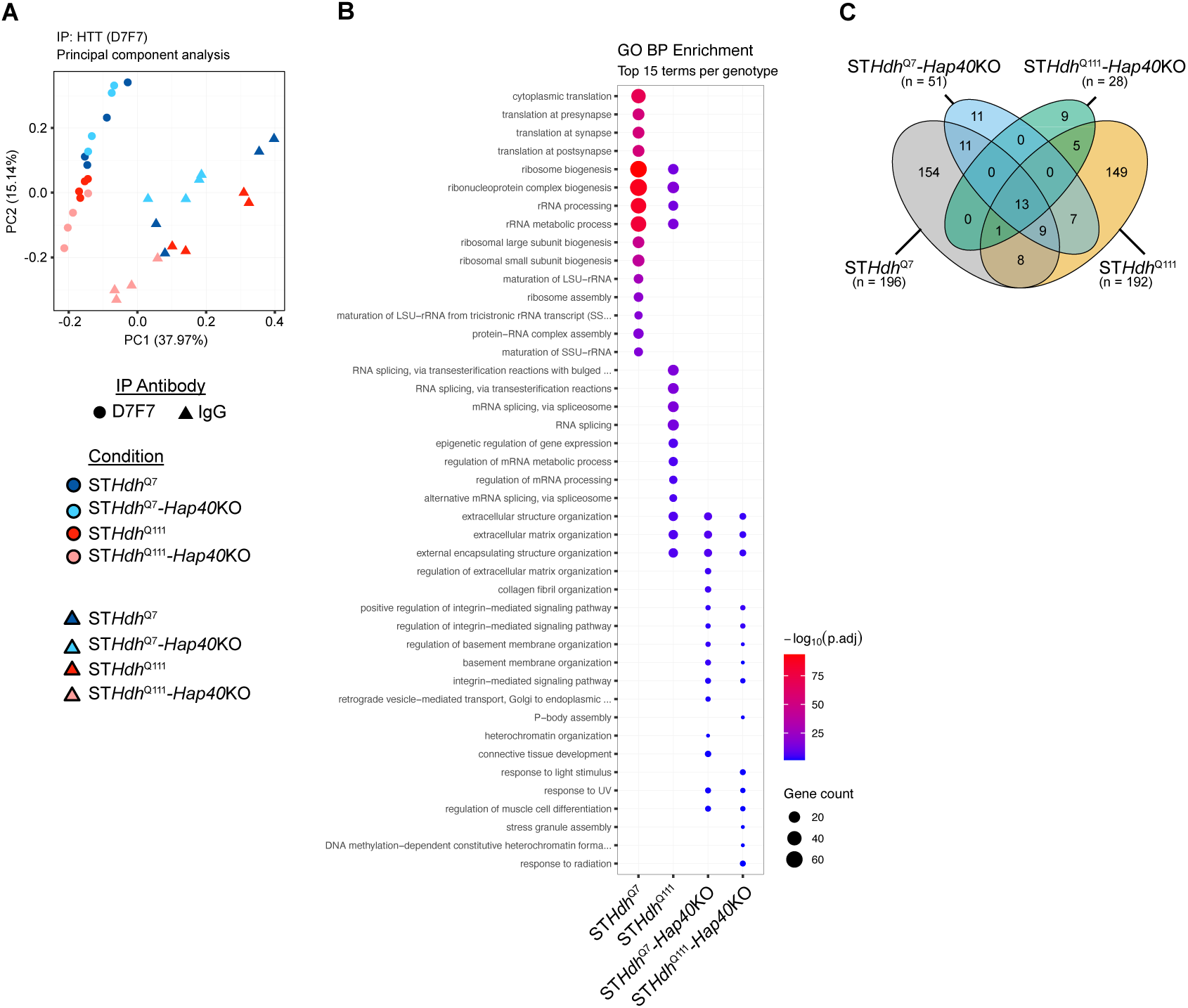
HTT-IP-MS analysis of striatal cells across wild-type, mutant HTT, and HAP40KO genotypes. **A)** Principal Component Analysis of the striatal Co-IP data. The PCA plot visualizes the proteomic variance across 32 samples based on 1,480 proteins identified. The first principal component (PC1) explains 32.15% of the total variance and primarily separates specific pull-downs (D7F7, circles) from non-specific controls (IgG, triangles). The second principal component (PC2) explains 13.4% of the variance and reflects differences related to the HTT genotype (Q111 vs. Q7) and HAP40 status. Biological replicates of the D7F7 groups cluster closely, indicating experimental reproducibility for the specific enrichment. In contrast, the IgG control samples show higher dispersion, reflecting the inherent stochastic nature of non-specific background noise. **B)** Gene Ontology enrichment analysis of the HTT interactome. The dot plot displays the top 15 significantly enriched Biological Process terms identified for each genotype. The size of the dots represents the number of proteins (Gene count) associated with each term, providing an estimate of the biological weight of the process. The color intensity represents the statistical significance as -log10 (p adjusted), with higher values (red) indicating stronger enrichment. **C)** Comparison of HTT protein interactors across genotypes. The Venn diagram illustrates the distribution of proteins significantly enriched in D7F7-immunoprecipitates compared to their respective IgG controls. Numbers indicate the count of proteins for each contrast. A core set of 13 proteins was identified across all four groups, representing stable HTT interaction partners independent of polyQ length or HAP40 presence.

**Figure S7:**
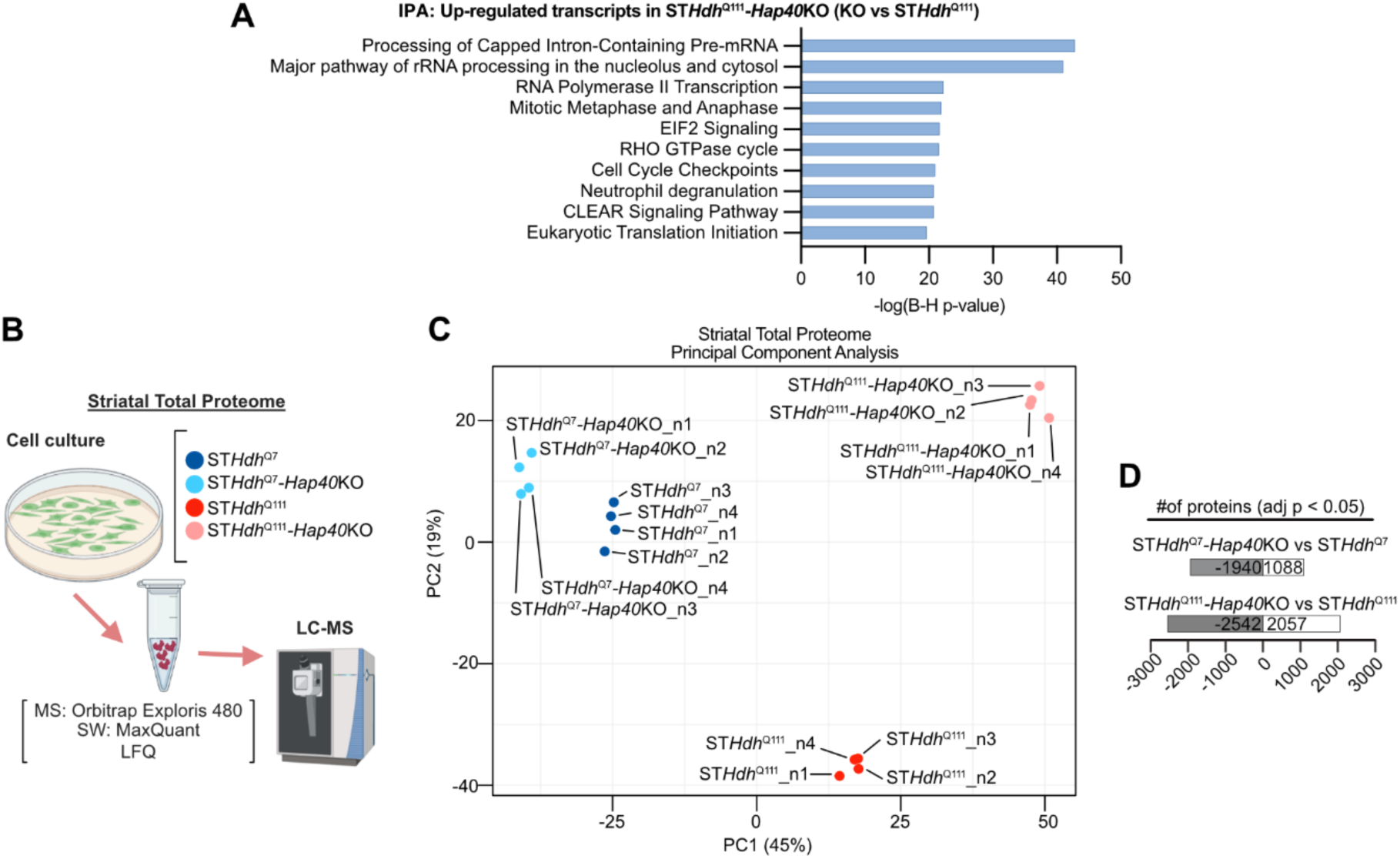
Analysis of transcriptome and proteome datasets of ST*Hap40*KO cell lines and controls. **A)** Significantly upregulated transcripts (adjusted p ≤ 0.05) in ST*Hdh*^Q111^-*Hap40*KO versus ST*Hdh*^Q111^ cells were analyzed by IPA to identify enriched canonical pathways, ranked by Benjamini–Hochberg FDR-adjusted p-values. **B)** Overview of the mouse striatal total proteome workflow. Four mouse striatal cell lines (ST*Hdh*^Q7^, ST*Hdh*^Q7^-*Hap40*KO, ST*Hdh*^Q111^, ST*Hdh*^Q111^-*Hap40*KO) were subjected to label-free quantitative (LFQ) LC-MS/MS. Four independent biological replicates were prepared for each genotype. Created in BioRender. Wanker, E. (2026) https://BioRender.com/ci9qwne **C)** Principal component analysis of striatal total proteome dataset. Points represent individual biological samples, colored by experimental group. The axes display PC1 and PC2. **D)** Counts of significantly differentially regulated proteins per contrast at the symbol accession level. Bars indicate up-regulated (log₂FC > 0) and down-regulated (log₂FC < 0) entries. Significance: adjusted p < 0.05 (Benjamini–Hochberg FDR).

**Figure S8:**
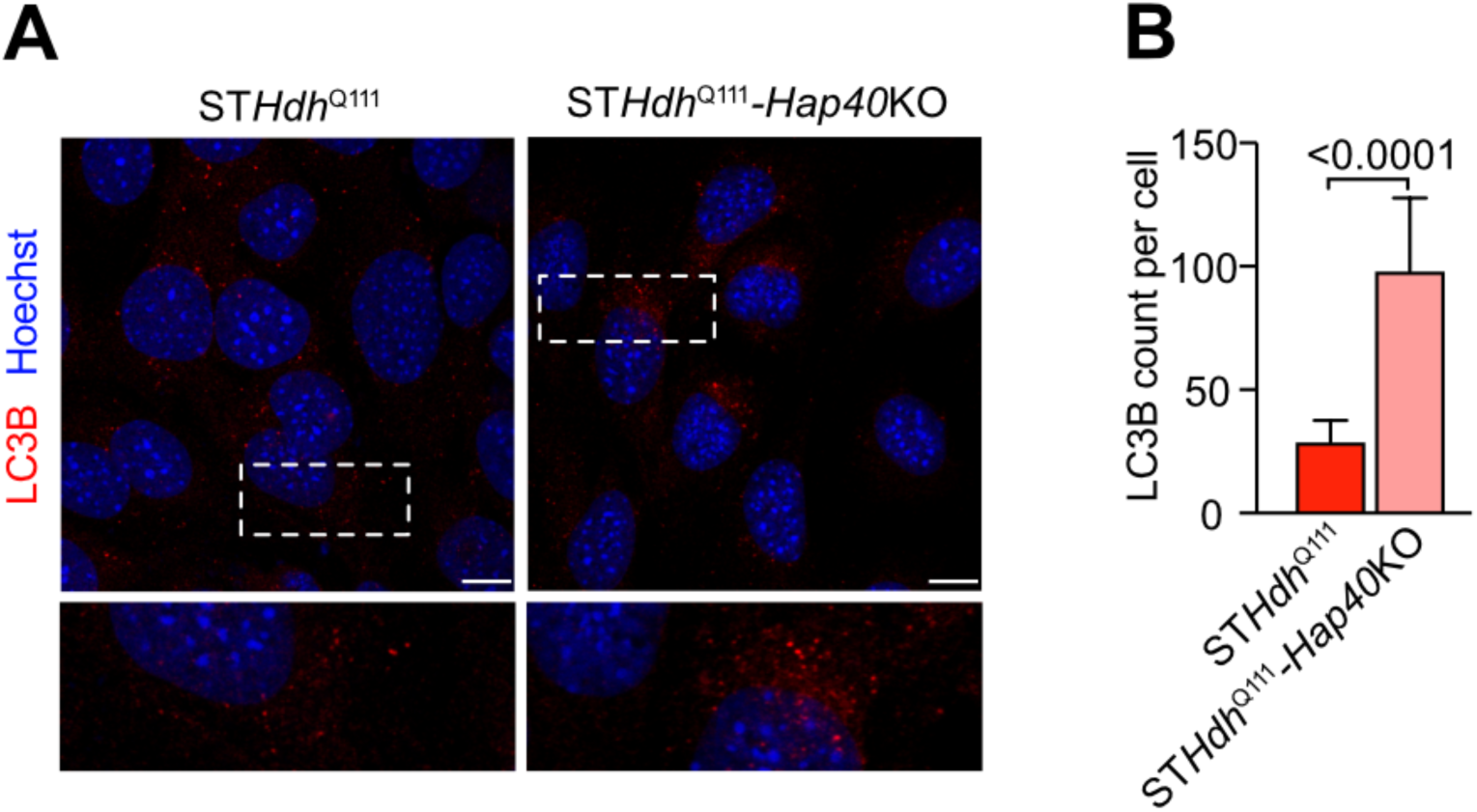
Assessment of autophagosome markers across HTT and HAP40 genotypes in striatal cells. **A)** Representative confocal images of striatal ST*Hdh*^Q111^ and ST*Hdh*^Q111^-*Hap40*KO cell lines immunostained for LC3B. Insets display magnified LC3B puncta for each genotype. Scale bar represents 10 µm. **B)** Quantification of LC3B count per cell. Twenty cells per genotype were analyzed across three independent biological replicates; data are shown as mean ± SD. Statistical significance was assessed by one-way ANOVA with Tukey’s multiple comparison test.

## Appendix

**Supplementary Table 1 (separate file).** iBAQ-based proteomic data from IP-MS experiments identifying HTTQ23-associated proteins in HEK293 WT, HEK293-HAP40KO, and HEK293-HTTKO cells.

**Supplementary Table 2 (separate file).** TMT-based total proteome profiles of HEK293 WT, HEK293-HAP40KO, and HEK293-HTTKO cells.

**Supplementary Table 3 (separate file).** Transcriptome profiles of striatal cell lines

**Supplementary Table 4 (separate file).** Total proteome profiles of striatal cell lines

